# β-catenin controls Afadin cortical recruitment to regulate planar epithelial cell division

**DOI:** 10.1101/2025.10.23.684278

**Authors:** Yue Meng, Pratchaya Rukthanapitak, Samantha Shimin Chang, Christine Siok Lan Low, Keng-Hui Lin, Pakorn Kanchanawong

## Abstract

Mitotic spindle orientation is tightly controlled in epithelia to preserve polarized tissue architecture. Here, we uncover a previously unrecognized role of the adherens junction protein β-catenin in regulating planar mitotic spindle positioning required for symmetric epithelial cell division. Using CRISPR/Cas9-mediated genome editing in MDCK cells, we found that β-catenin—but not its close homolog γ-catenin (plakoglobin)—is required for proper spindle orientation. Loss of β-catenin disrupts astral microtubule anchorage and impairs the localization of LGN–NuMA spindle orientation machinery. Mechanistically, β-catenin mediates spindle regulation via its N-terminal domain, independently of α-catenin binding. We showed that the cortical recruitment of Afadin during mitosis depends on β-catenin, but surprisingly, overexpression of either Afadin or ZO-1 rescues the spindle orientation in β-catenin deficient cells by restoring cortical LGN. Afadin depletion or dominant-negative Afadin abolished ZO-1-mediated compensation, thereby establishing Afadin as the central cortical integrator, potentiated by β-catenin, and operating through mass-action to recruit cortical LGN. Together, our findings define β-catenin as an upstream regulator of the Afadin-based cortical-spindle linkage that coordinates the crosstalk between adherens junctions and mitotic geometry to ensure epithelial homeostasis.

## Introduction

The formation of polarized epithelia is central to the evolution of animal multicellularity. Essential physiological processes such as nutrient absorption, barrier formation, and directional secretion, depend critically on epithelial polarization along the apicobasal axis, while dysregulation of epithelial polarity is associated with developmental disorders and tumorigenesis (Buckley and St Johnston, 2022; Vasioukhin et al., 2001; Zhan et al., 2008). Proper mitotic spindle orientation is an essential step in the establishment and maintenance of epithelial polarity and tissue architecture (Tang et al., 2018). For example, in-plane alignment of the spindle is critical for epithelial integrity during tissue expansion (Baer, Chanut-Delalande and Affolter, 2009). Conversely, asymmetric cell division involves the perpendicular alignment of the mitotic spindle poles and is requisite for differentiation (Speicher et al., 2008). In planar epithelial mitosis, crosstalk with the epithelial cell-cell junctions is known to provide positional cues for the mitotic spindle alignment (Gloerich et al., 2017; Tuncay et al., 2015). For instance, earlier studies have shown that E-cadherin contributes to spindle orientation via the recruitment of LGN, a conserved component in the mitotic spindle orientation machinery (Gloerich et al., 2017; Hart et al., 2017; Wang et al., 2018). However, complete ablation of E-cadherin results in surprisingly robust spindle orientation in 3D culture model (Otani et al., 2019). This suggests that AJ–spindle crosstalk may also involve additional AJ components whose roles remain incompletely defined.

Mitotic spindle orientation is governed by an intricate interplay between intracellular forces and membrane polarity complexes (Pannekoek, de Rooij and Gloerich, 2019). Motor proteins such as dynein and kinesin generate forces on astral microtubules to position the spindle (Kotak, Busso and Gonczy, 2012; Laan et al., 2012). Dynein, in particular, requires stable cortical anchoring to exert pulling forces. The evolutionarily conserved ternary complex of NuMA-LGN-Gαi is generally thought to play a central role (Chiu et al., 2016; Fankhaenel et al., 2023; Kiyomitsu and Boerner, 2021) interfacing between the force-generation machinery and the polarity cues encoded by cortical protein assemblies such as the Par complex (aPKC-Par3-Par6) (Durgan et al., 2011; Golub et al., 2017; Hao et al., 2010). Furthermore, LGN is known to interact with various polarity cues at the cell cortex to pre-specify spindle orientation prior to NuMA release upon Nuclear Envelope Breakdown (NEBD). For example, studies in MDCK cells revealed that LGN is excluded from the apical cortex through apical aPKC-mediated phosphorylation, with its subsequent localization at the lateral cortex hence favouring a planar orientation of the mitotic spindle (Hao et al., 2010), while in MCF10A breast epithelial cell model, Annexin A1 mediates LGN cortical localization to ensure in-plane spindle orientation (Fankhaenel et al., 2023). AJ-associated proteins such as E-cadherin or Afadin have also been shown to interact with the TPR domain of LGN (Carminati et al., 2016; Hart et al., 2017). Whether additional junction proteins may also interact with LGN or other spindle orientation components to mediate AJ-spindle crosstalk has not been fully elucidated, however.

Intercellular ligation at AJs depends on homophilic interaction of cadherin extracellular domains from opposing cells (Brasch et al., 2018; Charras and Yap, 2018; Takeichi, 2014). These are internally buttressed by cytoplasmic binding to Armadillo repeat-containing catenins such as p120-catenin, β-catenin, and γ-catenin (plakoglobin), which play multifaceted roles in AJ mechano-regulation and signalling. Notably, p120-catenin binds the juxtamembrane domain (JMD) of E-cadherin, stabilizing its plasma membrane localization by preventing endocytosis (Nanes et al., 2012). Distal to the JMD, β-catenin binds to E-cadherin and links to α-catenin or vinculin, forming mechanosensitive connections to the actin cytoskeleton (Bertocchi et al., 2017; Buckley et al., 2014; Morales-Camilo et al., 2024; Ozawa, Baribault and Kemler, 1989; Yonemura et al., 2010). Interestingly, the structural model of E-cadherin cytoplasmic domain complexing with LGN-TPR is reminiscent of its interaction with p120-catenin or β-catenin(Gloerich et al., 2017). Previous analysis appears to preclude p120-catenin from a role in spindle orientation(Yu et al., 2016), but whether β-catenin may contribute to AJ–spindle crosstalk has not been resolved. Beyond its role as a structural component of cadherin-based adhesions, β-catenin also functions as a transcriptional co-activator of TCF/LEF target genes in the Wnt signaling pathway(Clevers, 2006; Koelman, Yeste-Vazquez and Grossmann, 2022). Notably, aspects of β-catenin involvement in mitotic spindle apparatus have been reported previously. For example, studies in mouse embryonic fibroblasts (MEF) cells showed that phospho-β-catenin (Ser33/37/Thr41) is modified by conductin/axin2 and helps stabilize centrosome cohesion by interacting with centriole-associated component C-Nap1 (Hadjihannas, Bruckner and Behrens, 2010). Also, β-catenin has been found as Nek2 substrate during centrosome separation via Rootletin/C-Nap1 (Bahmanyar et al., 2008). However, whether and how β-catenin may be involved in mitotic spindle orientation remains unclear.

In this work, we identify β-catenin as an essential regulator of symmetric in-plane mitotic spindle orientation in the MDCK epithelial cell model. CRISPR/Cas9-mediated ablation of β-catenin, but not γ-catenin (plakoglobin), its close homologue, results in spindle orientation defects. Through re-expression of chimeric and truncation constructs, we show that the β-catenin N-terminal domain contributes to spindle orientation independently of α-catenin binding. β-catenin is required for stable LGN–NuMA localization at the lateral cortex during mitosis; its loss disrupts cortical astral microtubule organization. Furthermore, β-catenin is required for robust cortical recruitment of Afadin, but interestingly the overexpression of either Afadin or ZO-1 is able to rescue spindle orientation defects in β-catenin-deficient cells. Such compensatory effect depends on Afadin-ZO-1 interaction, suggesting that the Afadin-based cortical assembly may act through a mass-action mechanism to regulate cortical LGN. Consistent with this, β-catenin knockout MDCK fails to form single-lumen cysts in 3D Matrigel, but this defect is rescued upon ZO-1overexpression. Taken together, our findings delineate a previously unknown mechanism by which β-catenin functions as an upstream regulator of the cortical spindle-orientation machinery, coordinating cell-cell junction scaffolds with polarity cues to ensure robust symmetric cell division in epithelia.

## Results

### β-catenin but not γ-catenin (plakoglobin) is required for proper planar spindle orientation in MDCK

To investigate whether β-catenin may contribute to cell division and spindle orientation, we applied CRISPR/Cas9 genome editing(Bauer, Canver and Orkin, 2015) to ablate β-catenin encoding gene (gene name: CTNNB1) in MDCK II cells (**Supplementary Figure 1A-B**). Confocal microscopy of 2D monolayers reveals that β-catenin KO cells exhibit mitotic spindle orientation defects (**Figure 1A**). The spindle orientation angle relative to the horizontal plane was then quantified using immuno-stained γ-tubulin and α-tubulin as markers for spindle poles and spindle microtubules, respectively. As shown in **Figure 1B and 1D**, a significant elevation in spindle angle is observed in β-catenin KO compared to wild type cells, indicative of defective mitotic spindle orientation. Upon rescue by ectopic expression of full-length mouse β-catenin-mEmerald (**Supplementary Figure 1C**), in-plane orientation of the spindle is restored, thus corroborating the role of β-catenin in regulating spindle orientation.

**Figure 1.**
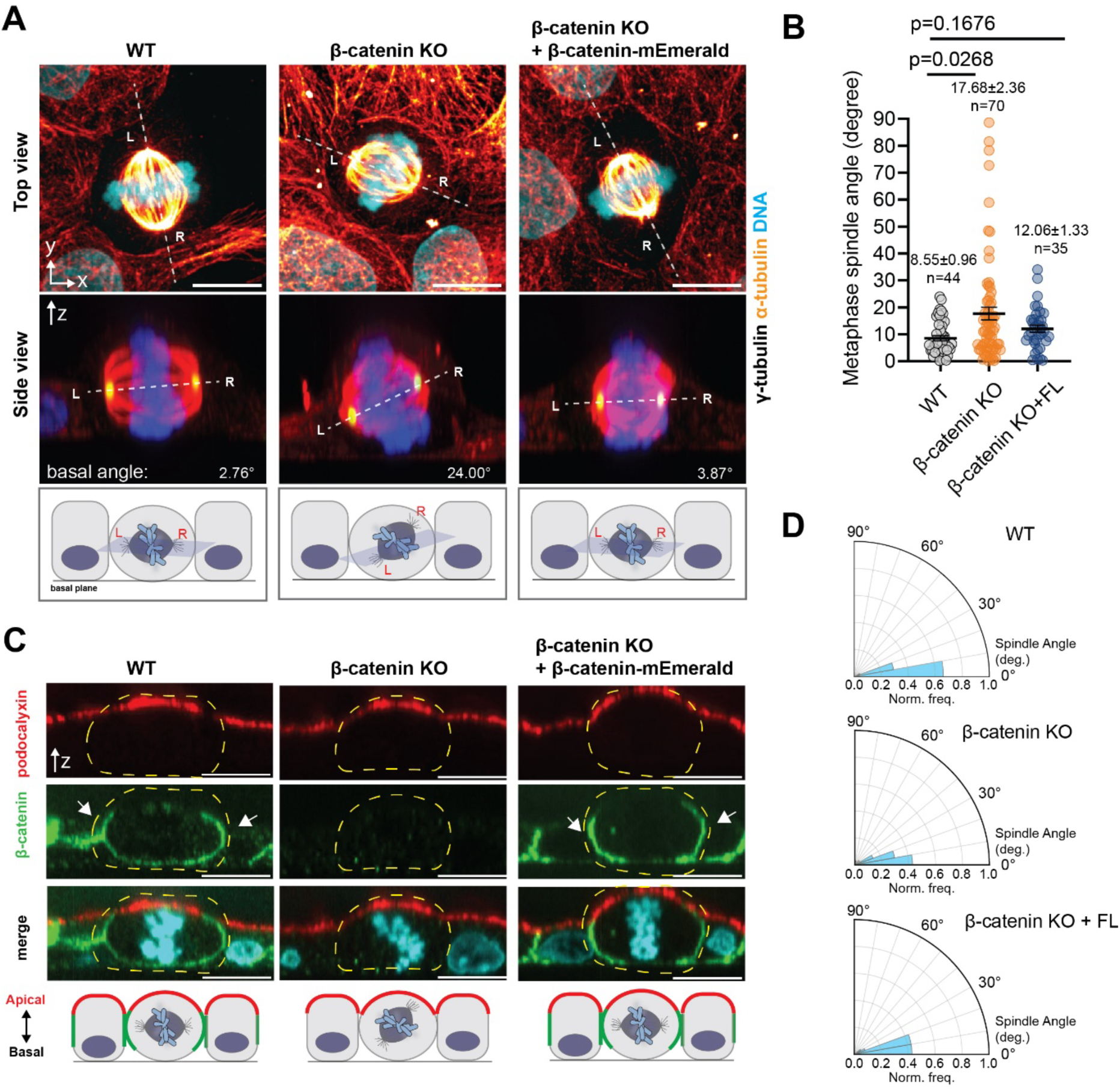
β-catenin is essential for planar epithelial cell division. (**A**) Composite multi-color confocal fluorescence micrographs of dividing MDCK cells, showing top view (upper row), side view (center row, zoomed side-view projection along the dashed lines in upper row), and schematic diagram of spindle orientation (bottom row). Wild-type (WT) (left) is compared with β-catenin KO (center) or β-catenin KO with β-catenin-mEmerald (FL) rescue (right). Top and side views are maximum intensity projection. Individual spindle poles are denoted L or R, respectively. Spindle angle corresponds to the angle between the basal plane and the spindle axis (white dashed line, center row). α-tubulin (red), γ-tubulin (green), DNA (cyan). (**B, D**) Quantification of metaphase spindle orientation angle distribution (B) and radial polar graph (**D**). Mean ± s.e.m., sample sizes, and p-value (Kruskal-Wallis) indicated on graph. Bin size (10 °) (D). (**C**) Side view confocal projection of dividing cells (dashed yellow lines) in epithelial monolayer, showing apical localization of podocalyxin (red), basolateral localization of β-catenin (green), as well as DNA (cyan). White arrows denote lateral cell-cell junction’s interface. Scale bars: 10 μm (A, C).

Since β-catenin is homologous with γ-catenin (plakoglobin), with overlapping functions reported at the cell-cell junctions (Solanas et al., 2016), we also used CRISPR/Cas9 to ablate the γ-catenin gene (gene name: JUP). Interestingly, in γ-catenin KO the mitotic spindle orientation angle remains similar to control (**Supplementary Figure 1D**). We therefore conclude that β-catenin but not γ-catenin (plakoglobin) is required for proper mitotic spindle orientation in MDCK II cells. As shown in **Supplementary Figure 1E**, single KOs of either β-catenin or γ-catenin are still able to form 2D epithelial monolayers, while the β-catenin/γ-catenin double KO is no longer able to form stable cell-cell junctions. Thus, while β-catenin and γ-catenin may have redundant functions in AJ integrity and epithelial monolayer formation, the role in ensuring in-plane mitotic spindle orientation appears to be unique to β-catenin.

To further understand the distinctive role of β-catenin in mitosis, we used 3D confocal microscopy to characterize β-catenin localization in dividing cells within contiguous 2D monolayer (side-view shown in **Figure 1C**). We found that in both dividing cells and non-dividing WT cells, β-catenin is clearly partitioned into the basolateral membrane and excluded from the apical membranes, as marked by podocalyxin (gp135), as expected. In β-catenin KO cells, the apical membrane appears to remain polarized, thus indicating that the apicobasal polarization of the plasma membrane is still retained despite β-catenin ablation.

During mitosis, astral microtubules emanating from the centrosomes are anchored in the cell cortex to orient and position the spindle (di Pietro, Echard and Morin, 2016; Mora-Bermudez, Matsuzaki and Huttner, 2014). Therefore, we next assess whether β-catenin may mediate the cortical attachment of astral microtubules. From confocal imaging with α-tubulin immunostaining, we observed a robust presence of astral microtubules in control cells. From live-cell tracking of EB1 comets, we also verified that microtubule dynamics were not significantly altered in β-catenin KO (**Supplementary Movies 1-3, Supplementary Figure 1F-G**). In contrast, astral microtubules are significantly depleted in β-catenin KO (Figure 2A), while being partially rescued upon the re-expression of full-length β-catenin, as quantified in **Figure 2B-C**. We further performed a comprehensive analysis of mitotic spindle and chromatin morphometric profiles, using Spindle3D software (Kletter et al., 2022). As shown in **Figure 2D-E**, this revealed that in β-catenin KO, chromatin dilation, chromatin volume, metaphase plate length, and metaphase plate width are significantly increased, in comparison to control cells or β-catenin rescue. In contrast, the spindle length, spindle width, and spindle aspect ratio remained similar to control (**Figure 2F**). Altogether, these results suggest that in β-catenin KO cells, astral microtubules likely become prone to detachment from the cortex or become otherwise destabilized. Consequently, the absence or mis-regulation of spindle pulling force relative to the cortex could give rise to the observed spindle mis-orientation (Anjur-Dietrich et al., 2024; Tame et al., 2016) and possibly aspects of chromosome congression. However, the global apicobasal polarization of the cells as well as the assembly and self-organization of the mitotic spindles appears minimally perturbed by the loss of β-catenin.

**Figure 2.**
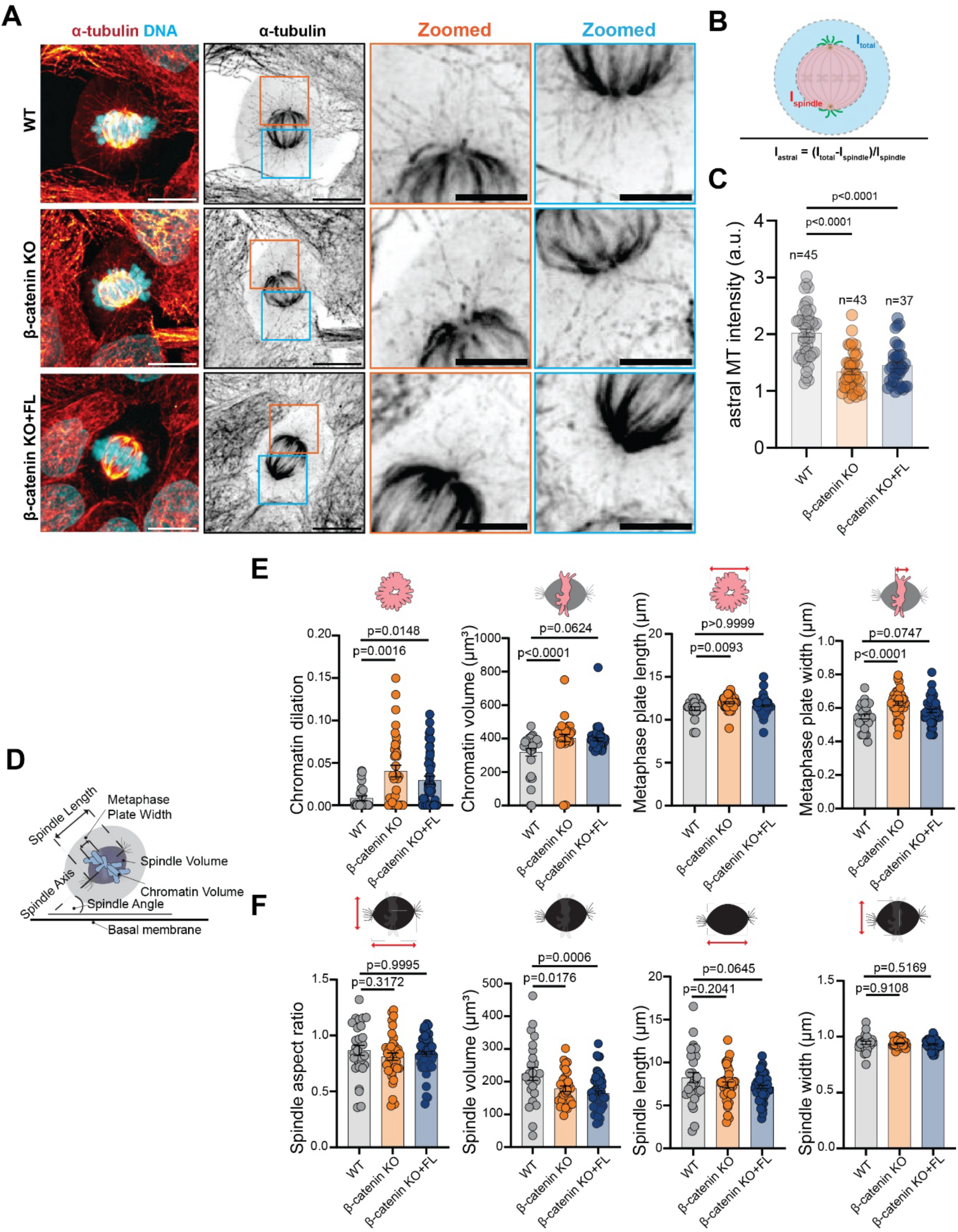
Effects of β-catenin loss on astral microtubules and mitotic spindle profiles. (**A**) Confocal fluorescence micrographs of microtubules and chromosomes in metaphase cells. (Left) Maximum intensity projection 2-color composite (tubulin, DNA), (inner left) α-tubulin channel, (inner right and rightmost) zoomed views corresponding to spindle poles denoted by orange or blue boxes in inner left column. (**B**) Schematic diagram for the quantification of astral microtubules density. (**C**) Quantification of astral microtubule density. Sample sizes and p-values indicated on graph. (**D**) Schematic diagram depicting spindle properties being analyzed. (**E, F**) Bar and swarm plots of the spatial properties of the metaphase chromosome (E) and the spindle (F). Sample sizes = 25 (WT), 36 (β-catenin KO), 45 (β-catenin KO+FL rescue). p-value indicated is from Kruskal-Wallis Test. Scale bars: 10 μm (A, left), 1 μm (zoomed insets).

### N-terminal domain of β-catenin is required for spindle orientation, independent of interaction with α-catenin

To delineate the molecular mechanisms by which β-catenin regulate spindle orientation, we next generated β-catenin constructs (**Figure 3A**) containing the truncation of either the N-terminal (1-124aa) or the C-terminal domain (669-745aa). The central armadillo repeats (ARM) domain was retained in the truncation constructs due to its indispensable role as the E-cadherin binding site for membrane docking. As shown in **Figure 3A**, when expressed in β-catenin KO cells, both truncation constructs are capable of localization to the cell-cell junctions. Since full-length β-catenin is able to rescue spindle mis-orientation (**Figure 1A**), we assayed the ability of the truncation constructs to rescue spindle orientation. From quantification of the spindle orientation, we found that the C-terminal truncation (β-catenin ΔC construct) significantly rescue spindle orientation defect (**Figure 3B**). In contrast, spindle mis-orientation remains prevalent in KO cells expressing N-terminal truncated β-catenin. Amino acid sequence alignment shows that β-catenin and γ-catenin share nearly 83% amino acids similarity in ARM domain but only 57% in N-terminal and 15% in C-terminal domains(Solanas et al., 2016). While sequence homology implies that the C-terminal domains are more likely to contribute to their distinct functions, our results whereby the C-terminal domain truncation of β-catenin can rescue proper spindle orientation instead, indicate that the unique determinant of β-catenin mitotic function may reside in the N-terminal region (**Figure 3A-B**). Corroborating this, we found that a chimeric construct in which the N-terminal domain (1-124aa) of β-catenin is replaced by the N-terminal domain of γ-catenin (1-134aa) was unable to rescue spindle mis-orientation, while a chimeric construct in which the N-terminal domain (1-124aa) of β-catenin fused with γ-catenin ARM domain and C-terminal domain can restore planar spindle orientation. Therefore, we conclude that the N-terminal domain of β-catenin is required for proper in-plane mitotic spindle orientation.

**Figure 3.**
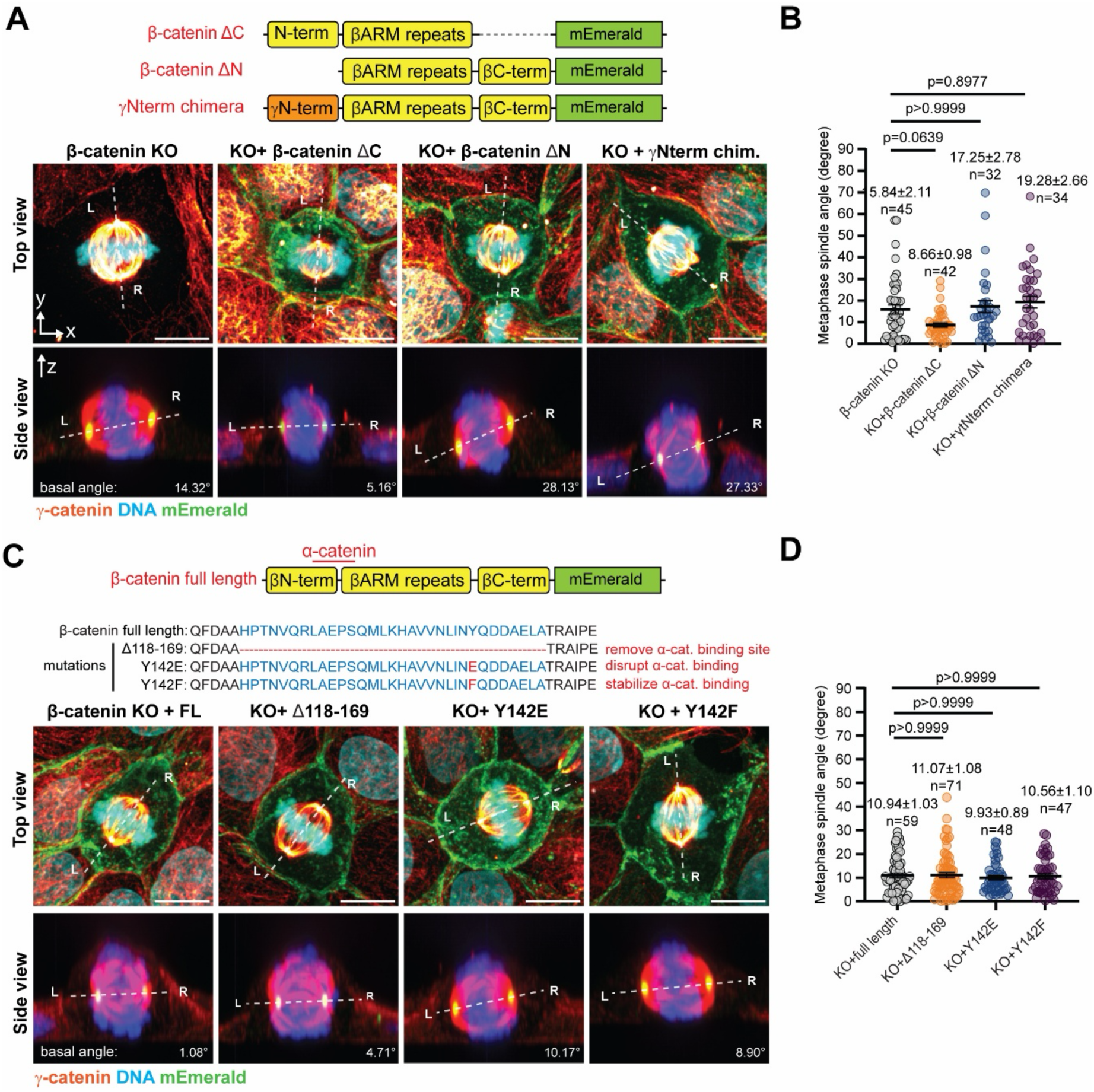
β-catenin mediates spindle orientation through its N-terminal domain independent of α-catenin. (**A, C**) Schematic of truncation, chimeric, or mutant constructs (top), and representative confocal images of dividing β-catenin KO cells expressing each construct (bottom). Sideview panels correspond to projection along the dotted lines in top panel, with spindle axis and spindle angle relative to the basal plane indicated. (**B, D**) Swarm plots of metaphase spindle orientation angle. Mean ± s.e.m., sample sizes, and p-value (Kruskal-Wallis) indicated on graph. Scale bars: 10 μm (A, C).

The N-terminal domain of β-catenin overlaps with the binding site for α-catenin (residues 118-169 (Pokutta and Weis, 2000)), a major AJ component that provides direct mechanosensitive linkage to the actin cytoskeleton (Pokutta and Weis, 2000; Yonemura et al., 2010). Hence, we wondered whether α-catenin may also contribute to β-catenin-mediated spindle orientation. To test this, we generated β-catenin constructs with Δ118-169 deletion. Additionally, since residue Y142 on β-catenin has been shown to be vital for the maintenance of α-catenin binding and junctional mechanical property (Le, Yu and Yan, 2019), we also generated the Y142E and Y142F point mutants that respectively disrupt α-catenin interaction or stabilize α-catenin interaction (**Figure 3C-D**). However, upon the expression of these constructs in β-catenin KO cells, we found that they are all capable of restoring proper spindle orientation, with no significant differences in the spindle orientation angle compared to full-length control. These results indicated that α-catenin binding minimally contribute to spindle orientation, arguing against its role in mediating β-catenin-dependent spindle regulation. Note that since endogenous γ-catenin is still expressed in the β-catenin KO cells, α-catenin recruitment by γ-catenin may potentially compensate for the loss of β-catenin. However, this possibility is discounted by the inability of the chimeric N-terminal-γ-catenin::β-catenin construct to rescue spindle orientation defect (**Figure 3A-B**). A systematic truncation and mutagenesis analysis to identify key sequence motifs in the N-terminal domain along with salient binding partners will be an important direction for future work to reveal additional regulators of β-catenin-dependent spindle orientation.

### β-catenin stabilizes LGN at the lateral cortex during mitosis

In addition to cell-cell junction localization, β-catenin undergoes nuclear/cytoplasmic shuttling as part of the Wnt signalling pathway (Nusse and Clevers, 2017). Nuclear import of β-catenin is regulated by the phosphorylation of residue S675 in the Armadillo domain by p21-Activated kinase 4 (PAK4) (Chetty et al., 2020; Selamat et al., 2015). To explore whether the impairment of Wnt signaling may contribute to the observed spindle orientation defect, we pharmacologically inhibited PAK4 using PF-378309 (Murray et al., 2010). As shown in **Supplementary Figure 2A-B**, the efficacy of PAK4 inhibition is readily apparent from the observation of widespread lagging chromosomes due to the roles of PAK4 in mitotic spindle positioning and anchoring (He et al., 2019). However, spindle orientation appears to be unperturbed by PAK4 inhibition, suggesting that β-catenin and PAK4 probably contribute to mitosis via different pathways. Based on these results, thereafter we focused on the cortical functions of β-catenin.

Astral microtubules are anchored to the cortex via NuMA-LGN-Gαi and dynein-dynactin complexes (Sun et al., 2021). Previous study has shown that LGN interact with E-cadherin cytoplasmic tail via its TPR domain during initial recruitment (Gloerich et al., 2017; Hart et al., 2017). Since β-catenin localization to the cortex also depends on binding to E-cadherin cytoplasmic tail, we sought to investigate whether its mitotic spindle role may involve LGN by characterizing the endogenous LGN localization during cell cycle in WT cells. As shown in **Figure 4A and Supplementary Figure 3**, we observed that during interphase endogenous LGN are observed in cytosolic punctate structures. Upon entering prophase, LGN localization at the cell-cell junctions noticeably increases, becoming symmetrically distributed along the plasma membrane (asterisks, **Figure 4B, D**). As mitosis progresses, further accumulation of LGN at the spindle poles becomes pronounced at metaphase (**Figure 4D**). In WT, LGN localization at the plasma membrane persists during anaphase, while becoming more cytosolic upon cytokinesis (Du and Macara, 2004; Kaushik et al., 2003). In contrast, in β-catenin KO cells, we observed a marked reduction of LGN cortical intensity at metaphase. This can be restored to the control WT level upon the expression of full-length β-catenin-GFP as well as ZO-1-GFP, or Afadin-mCherry conditions as described further below (**Figure 4C**). Additionally, the distribution of cortex-associated LGN becomes asymmetric, instead of the symmetrical distribution observed in WT control (**Figure 4B, D and E**). This raises the possibility that the asymmetrical cortical localization of LGN may contribute to improper attachment of spindle machinery to the cortex in β-catenin KO, thereby giving rise to the observed spindle orientation defects.

**Figure 4.**
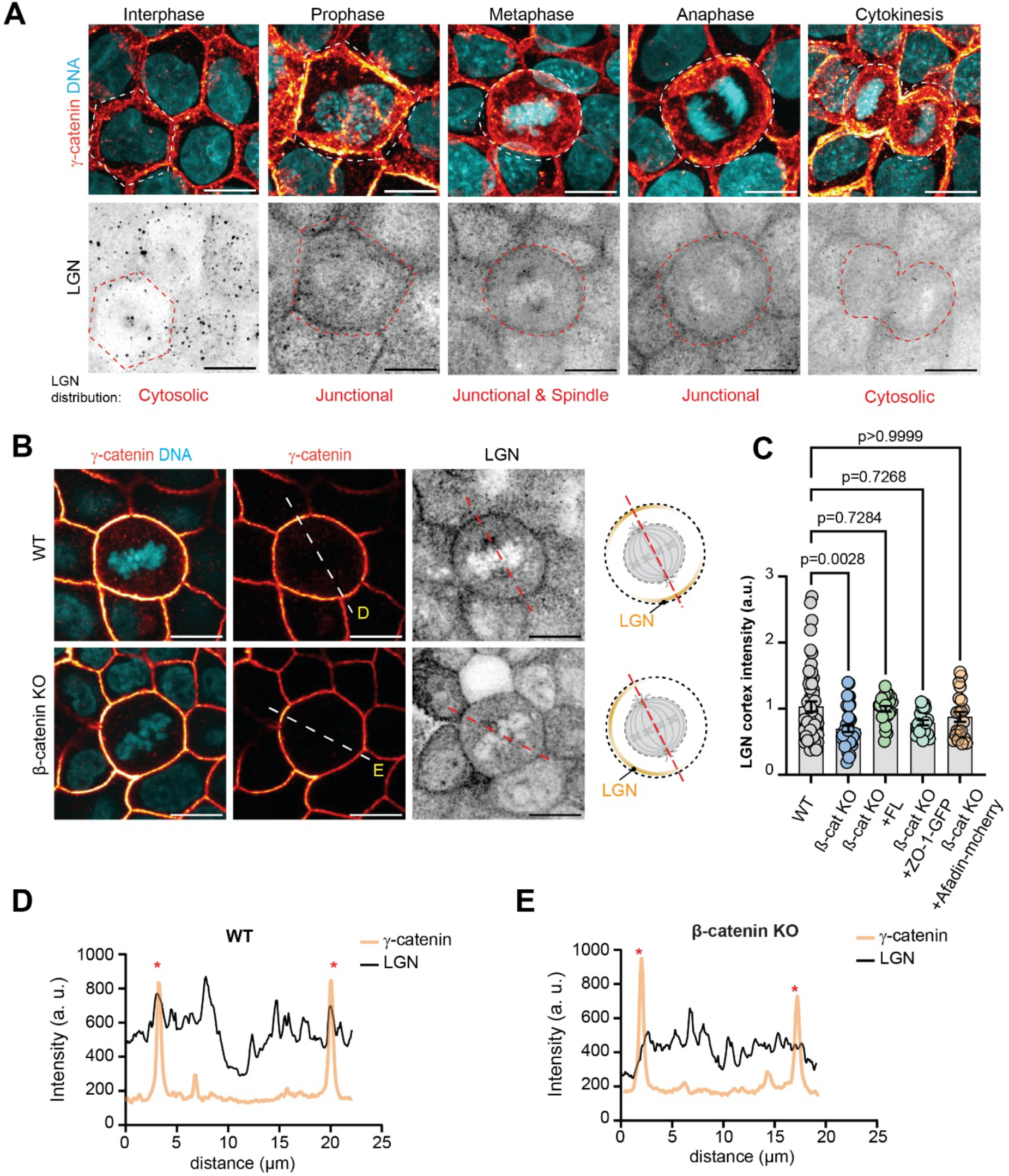
β-catenin stabilizes LGN cortical localization during mitosis. (**A**) Confocal immunofluorescence micrograph (Maximum Intensity Projection) of endogenous LGN, cell-cell junction markers γ-catenin, and chromosomes in WT, showing changes in LGN distribution at different cell cycle stages. (**B**) Confocal slice corresponding to mid-spindle z-plane comparing metaphase distribution of LGN between WT and β-catenin KO, which exhibit asymmetric LGN cortical distribution. (**C**) Swarm plots comparing the cortical intensity of endogenous LGN in WT, β-catenin KO, and β-catenin KO with expression of β-catenin full-length, ZO-1-GFP, or Afadin-mcherry constructs, respectively. (**D**, **E**) Intensity profiles along the spindle axis denoted in (**B**) comparing WT and β-catenin KO, showing decreased cortical LGN intensity (red asterisks) in β-catenin KO. Scale bar: 10 μm (A, B).

Subsequent to LGN, the recruitment of NuMA to the spindle orientation machinery has been shown to mediate spindle focus due to its C-terminal microtubule bundling activity (Hueschen et al., 2017). More specifically, NuMA recognizes the minus-ends of the microtubules and interact with the dynein-dynactin complex to enable the movement of NuMA-bound microtubules towards the centrosomes, resulting in the spindle focusing on the poles. The C-terminal domain of NuMA also defines its cortical localization during mitosis via its binding to the TPR domain of LGN. Therefore, we next investigate whether NuMA is affected by asymmetric LGN accumulation. As shown in **Figure 5A-B**, we characterize the localization of NuMA during cell division from interphase to cytokinesis, comparing WT with β-catenin KO. We observed that in WT during the interphase, NuMA is ensconced in the nucleus, consistent with previous studies (Chu et al., 2016). Following NEBD, NuMA disperses from the nucleus, with nascent accumulation near the centrosomes. During metaphase, NuMA robustly accumulates at the spindle poles. As anaphase proceeds toward telophase and cytokinesis, spindle-pole localized NuMA becomes progressively reduced in intensity, becoming largely cytosolic upon cytokinesis. In contrast, in β-catenin KO, we observed a delay in the accumulation of NuMA at spindle poles. Although NuMA localization at the spindle poles during metaphase are still clearly observed, NuMA appears to dissipate early in late metaphase and largely dissociates from the spindle poles as anaphase progresses. In parallel, NuMA spindle intensity at metaphase was significantly reduced in β-catenin KO cells (**Figure 5C**), indicating that NuMA recruitment to, or the stability at, the spindle apparatus is compromised in the absence of β-catenin. To assess their biochemical interaction, we also performed cell cycle synchronization to enrich mitotic cells followed by immunoblotting of β-catenin KO rescued with full-length β-catenin. To synchronize metaphase cells, RO3306 (arrest at G2/M border) and MG132 (to inhibit proteasome) was applied (Eli et al., 2025). Since β -catenin was recently shown to undergo liquid-liquid phase separation (Monster et al., 2025), co-immunoprecipitation with NuMA was performed using synchronized cells with or without treatment by the cell-permeable cross-linker dimethyl 3,3′-dithiobispropionimidate (DTBP) which facilitates the capture of weak interaction (Humphries et al., 2009). As shown in **Figure 5D**, we found that NuMA was co-immunoprecipitated with β-catenin both in the presence or absence of DTBP, indicative of robust biochemical interaction. Taken together, these results are consistent with our hypothesis that β-catenin is required for proper spindle apparatus stability through interaction with NuMA.

**Figure 5.**
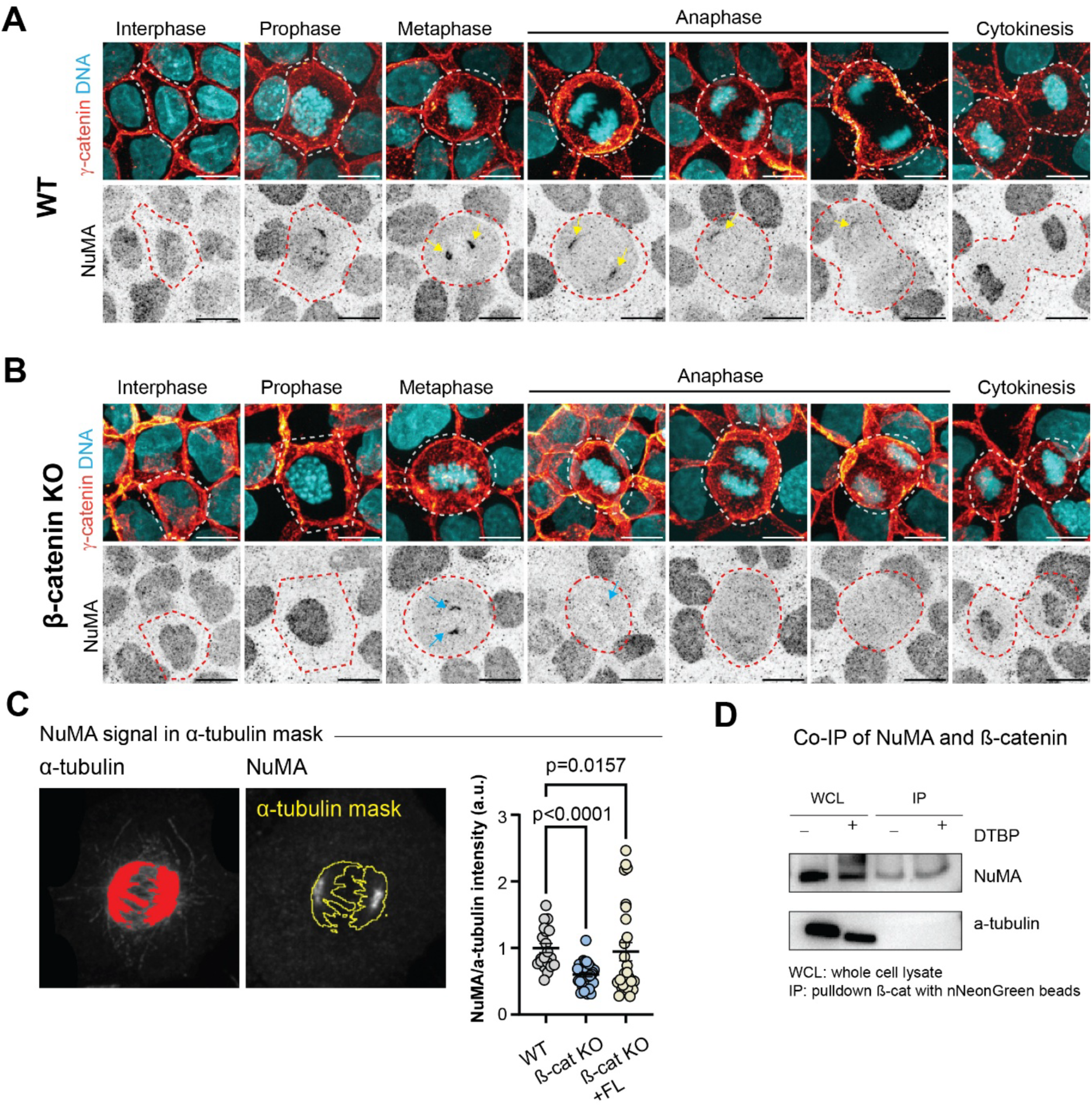
β-catenin stabilizes NuMA spindle pole localization during mitosis. Confocal immunofluorescence micrographs of endogenous NuMA, cell-cell junction marker γ-catenin, and chromosomes in WT (**A**) or β-catenin KO (**B**) at various cell cycle stages. In WT, NuMA is confined in the nucleus at interphase, released upon NEBD (prophase), and recruited to spindle poles (yellow arrows, metaphase to anaphase), before shuttling back to the nucleus upon cytokinesis. In β-catenin KO, NuMA recruitment to spindle poles appears to be delayed until metaphase (blue arrows) and diminished early in anaphase. (**C**) Comparison of NuMA signal quantified within α-tubulin mask between WT, β-catenin KO and β-catenin KO expressing β-catenin full-length. (**D**) Co-immunoprecipitation of NuMA and β-catenin-mEmerald expressed in β-catenin KO cells. Lysates were collected from synchronized cells, with or without DTBP treatment. Scale bars, 10 μm (A-B).

### Tripartite compensatory network of β-catenin/ZO-1/Afadin mediates planar mitotic spindle orientation

In order to monitor the cell-cell junctions during live imaging, we also made use of CRISPR/Cas9 to generate a β-catenin KO of a MDCK cell line with stable overexpression of ZO-1-EGFP and Histone H2B-mCherry described earlier (Saw et al., 2022) (**Supplementary Fig. 4A**). Surprisingly, we found that mitotic spindle orientation appears normal in β-catenin KO generated in this background (**Figure 6A-B**). This led us to test whether ZO-1 overexpression is capable of rescuing mitotic defect in the earlier β-catenin KO clones generated from parental MDCK II. As shown in **Supplementary Figure 4B**, we observed a similar rescue of in-plane spindle orientation. The ability of ZO-1 to rescue mitotic spindle defects in the absence of β-catenin implicates a previously unknown compensatory pathway that regulates mitotic spindle orientation, leading us to further characterize the underlying mechanism.

**Figure 6.**
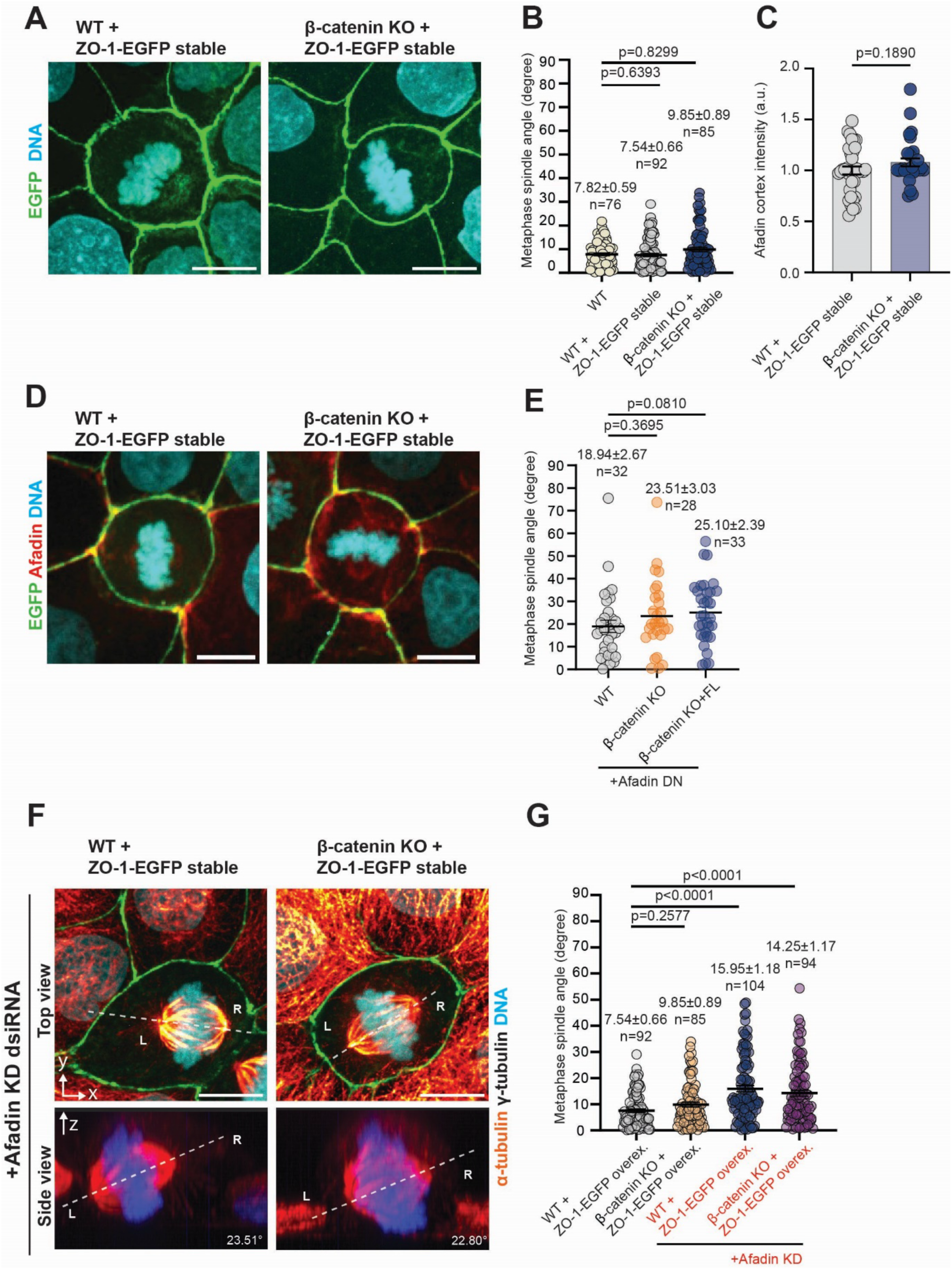
Compensation of β-catenin depletion by ZO-1 in planar spindle orientation. (**A**) ZO-1 overexpression compensates for β-catenin knockout misorientation defects. Confocal images of metaphase MDCK with ZO-1-EGFP (green)/Histone H2B (blue) stable expression comparing parental (WT) and β-catenin KO. (**B**) Quantification of metaphase spindle angles in WT, WT overexpressing ZO-1-EGFP, and β-catenin KO overexpressing ZO-1-EGFP. Sample sizes, mean ± s.e.m, and p-values indicated on graph. (**C-D**) ZO-1 overexpression restores cortical intensity of endogenous Afadin. (D) Confocal immunofluorescence of Afadin (red) in MDCK with ZO-1-EGFP/Histone H2B stable expression, with quantification shown in (C). (**E-G**) ZO-1-mediated compensation is dependent on Afadin. (E) Quantification of metaphase spindle angle, comparing MDCK WT, β-catenin KO, and KO rescued by β-catenin-mEmerald (FL) re-expression, with expression of Dominant-Negative (DN) Afadin. (F) Representative confocal images of dividing cells. WT or β-catenin KO in ZO-1-EGFP/Histone H2B stable background are treated with Afadin dsiRNA. Side view panels (bottom) correspond to projection along dashed lines in top panels. Spindle axis and angle relative to horizontal plane indicated. (G) Quantification of metaphase spindle orientation corresponding to conditions in (A) and (F). Sample sizes, mean ± s.e.m., and p-values indicated on graphs. Scale bars, 10 μm (A, D, F).

Our results so far have linked β-catenin with the cortical spindle orientation machinery, yet β-catenin contains no known direct binding sites with the cytoskeletons. Having discounted the role of α-catenin binding site to β-catenin earlier (**Figure 3**), we instead consider the role of ZO-1. Although ZO-1 has no known direct binding with β-catenin, a number of proteins can serve as intermediary including α-catenin and Afadin(Itoh et al., 1997). Previous studies in *Drosophila* neuroblasts and muscle progenitor cells have shown that the ablation of Canoe/Afadin prevents cortical accumulation of Mud/NuMA and disrupts the spindle alignment through disruption of the interaction with Pins/LGN(Speicher et al., 2008; Wee et al., 2011). Indeed, in mammalian epithelial cells, Afadin has been shown to bind to both LGN and F-actin directly and is required for correct spindle orientation (Carminati et al., 2016). Since Afadin could also localize to AJs with reported binding interactions to both actin cytoskeleton and β-catenin(Ikeda et al., 1999), we hypothesize that Afadin may also mediate β-catenin-dependent spindle orientation.

To test this, we first examined the localization of Afadin during metaphase. As shown in **Figure 7A-B**, in WT, Afadin exhibits clear localization at both the mitotic spindles and the plasma membrane. In contrast, in β-catenin KO, while Afadin localization to the spindle remains robust, its cortical localization is significantly reduced. Notably, upon the re-expression of full-length β-catenin, the cortical localization of Afadin is restored (**Figure 7A**), indicating that β-catenin is required for Afadin cortical targeting. To further validate the role of Afadin, we next knock down Afadin using dsiRNA (**Supplementary Figure 5A**). As shown in **Figure 7E-F**, we observed that Afadin KD in both WT and β-catenin full-length rescued cells result in defective spindle orientation as indicated by the spindle orientation quantification. In conjunction, a significant loss of astral microtubules similar to in β-catenin knockout cells compared to the control WT was also observed (**Supplementary Figure 5C-D**). Interestingly, when Afadin-mCherry was overexpressed in β-catenin KO cells, we found that proper spindle orientation is restored (**Figure 7C-D, Supplementary Figure 5B**), analogous to the rescue by ZO-1 overexpression. Taken together, these results corroborate the requirement of Afadin in spindle positioning and reveal the presence of a tripartite compensatory network consisting of β-catenin, ZO-1, and Afadin that mediate in-plane mitotic spindle orientation.

**Figure 7.**
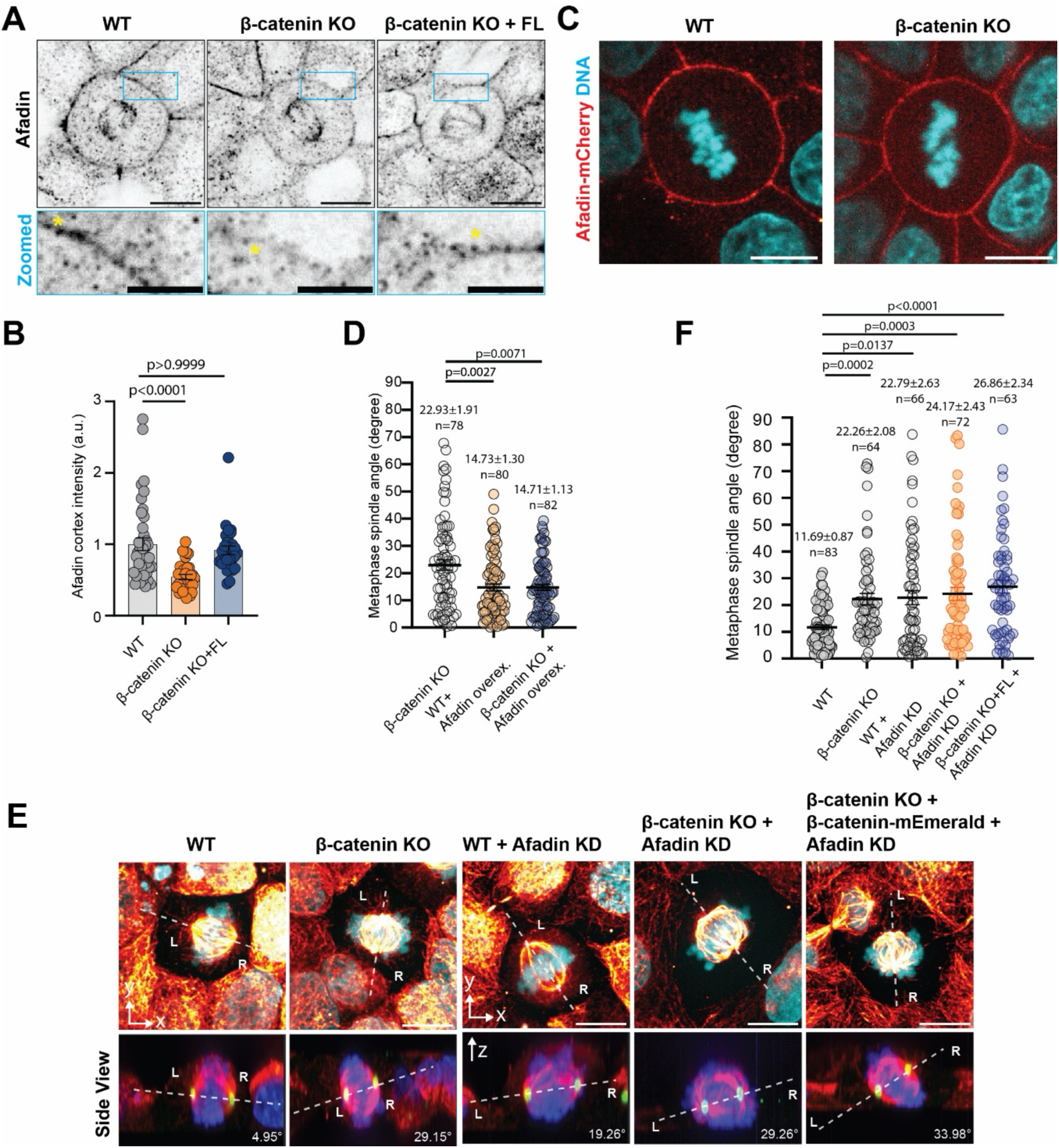
β-catenin acts upstream of Afadin in spindle orientation regulation. (**A**) Afadin localization to the cortex depends on β-catenin. Inverted contrast confocal immunofluorescence of endogenous Afadin (top row) and zoomed views corresponding to top row insets (bottom row), with cortex regions denoted by yellow asterisks. In metaphase β-catenin KO cells, Afadin retains spindle poles localization but is depleted from the cortex. (**B**) Quantification of cortical Afadin intensity, comparing WT, KO, and KO expression full-length (FL) β-catenin. Sample size: 40 (WT), 34 (KO), 29 (FL). (**C-D**) Overexpression of Afadin-mCherry compensates for β-catenin loss. Confocal slice of dividing WT or β-catenin KO overexpressing Afadin-mCherry (**C**) and metaphase spindle orientation quantification (**D**). (**E**) Depletion of Afadin disrupts planar spindle orientation. Representative confocal images of dividing cells with Afadin KD (top row). Sideview panels (bottom row) correspond to projection along the dotted lines in top panel, with spindle axis and spindle angle relative to the basal plane indicated. (**F**) Quantification of metaphase spindle angle in response to Afadin depletion. Mean ± s.e.m, sample sizes, and p-values indicated on graphs. Scale bar, 10 μm (A, C, E), 5μm (A, zoomed insets).

We next sought to better elucidate the functional hierarchy within such network. Thus far, either overexpression of Afadin or ZO-1 are able to rescue spindle orientation defects caused by the absence of β-catenin. Immunostaining of endogenous Afadin in ZO-1 overexpressing WT as well as β-catenin reveals comparable extent of Afadin cortical recruitment (**Figure 6C**), suggesting that overexpression of ZO-1 could compensate for the loss of β-catenin-dependent Afadin recruitment to the cortex, consistent with their known interactions (Beutel et al., 2019). Additionally, we observed that the junctional distribution of endogenous ZO-1 becomes fragmented in β-catenin KO cells, in contrast to the continuous distribution in WT (**Supplementary Figure 4C**), suggesting that β-catenin may also be required for proper ZO-1 localization at tight junctions. To distinguish the functional hierarchy between Afadin and ZO-1, we next performed dsiRNA-mediated KD of Afadin knockdown on the ZO-1 overexpression WT or β-catenin-KO background (**Supplementary Figure 5E**). This led to spindle orientation defects in both groups (**Figure 6F-G**), indicating that ZO-1 overexpression may be insufficient for compensating for Afadin loss. We corroborate this by using a dominant negative (DN) mutant of Afadin which abolishes ZO-1 interaction by S216A/S1090A mutations (Wu et al., 2020). Consistent with the Afadin KD results, the expression of Afadin-DN induces spindle orientation defects throughout WT, β-catenin-KO, and rescued groups (**Figure 6D-E, Supplementary Fig 5F**).

Subsequently we perform co-immunoprecipitation analysis to assess biochemical interactions between β-catenin, ZO-1, and Afadin. As shown in **Supplementary Figure 6A**, in cells overexpressing ZO-1-GFP, Afadin is pulled down with ZO-1 in both WT and β-catenin KO conditions, indicating that β-catenin is dispensable for the interaction between Afadin and ZO-1. Co-immunoprecipitation of Afadin and β-catenin was then performed in β-catenin KO MDCK cells rescued with a full-length β-catenin-mNeonGreen construct, with and without DTBP as both have been shown to localize in condensates (Monster et al., 2025). Notably, as shown in **Supplementary Figure 6B**, Afadin robustly co-immunoprecipitated with β-catenin regardless of crosslinking, consistent with earlier study and indicative of robust biochemical interactions. Similarly, ZO-1 was also observed to co-immunoprecipitate with β-catenin (**Supplementary Figure 6C**), consistent with previous evidence (Rajasekaran et al., 1996). Notably, unlike NuMA interaction with β-catenin (**Figure 5D**) which requires cell cycle synchronization to enrich mitotic cells, the interactions between ZO-1, Afadin, and β-catenin were detected in unsynchronized populations, suggestive of their presence as constitutive complexes. Collectively, these findings corroborate a functional hierarchy in which cortical Afadin serves as the primary link with the mitotic spindle orientation apparatus via LGN binding, while β-catenin and ZO-1 serve as mutually supportive scaffold for Afadin that localize it to the cortical complexes, thereby enabling robust spindle orientation.

### Requirement of β-catenin in single lumen formation during MDCK cystogenesis

Finally, to evaluate the role of β-catenin in regulating mitotic spindle orientation in 3D epithelial model, we took advantage of the ability of MDCK cells to form single-lumen cyst *de novo* when cultured in 3D Matrigel. MDCK cystogenesis involves two major checkpoints, with the first being the formation of the Apical Membrane Initiation Site (AMIS) (Bryant et al., 2010) which involved transcytosis of apical proteins (such as podocalyxin) from the extracellular membrane-facing surface to the nascent apical surface, thus defining the Pre-Apical Patch (PAP). This is subsequently followed by proper mitotic spindle orientation perpendicular to the apicobasal axis during lumen expansion phase (Lujan et al., 2017) (Martin-Belmonte et al., 2007; Martin-Belmonte et al., 2008). In 2D epithelial monolayer, aberrant mitotic spindle orientation occurs upon β-catenin ablation as described above. However, it is not known whether the earlier steps of AMIS and PAP formation are also affected.

We first categorized the cyst morphology formed by single-cell seeding in 3D Matrigel at 24-, 48-, and 72-hours post seeding, using immuno-staining of podocalyxin localization to define the apical membrane. In the canonical model (Bryant et al., 2010), at the earliest 2-cell stage, podocalyxin localization in WT cyst is confined to the ECM-facing surface, here denoted stage I. Subsequently, podocalyxin is transcytosed, appearing as internalized vesicles (denoted as stage II), before being finally exocytosed at the PAP which is located at the nascent luminal surface of the cyst, thus completing the inversion of the podocalyxin-containing membrane, forming proper apical domain (stage III). As shown in **Figure 8A-B**, we found that at 24 h in WT control, initial PAP formation is well under way with nearly equal distribution between stage I, II, and III morphology. Similarly, for β-catenin KO at 24h, nearly similar proportion of stage I, II, and III morphology are observed, indicative of progressive advance through PAP formation. Likewise, at 48h and 72h post-seeding, both WT and β-catenin KO progress at a similar pace, with most cysts able to complete the inversion of podocalyxin by 72h. Thus, we conclude that β-catenin is dispensable for AMIS formation, exocytosis, and PAP formation.

**Figure 8.**
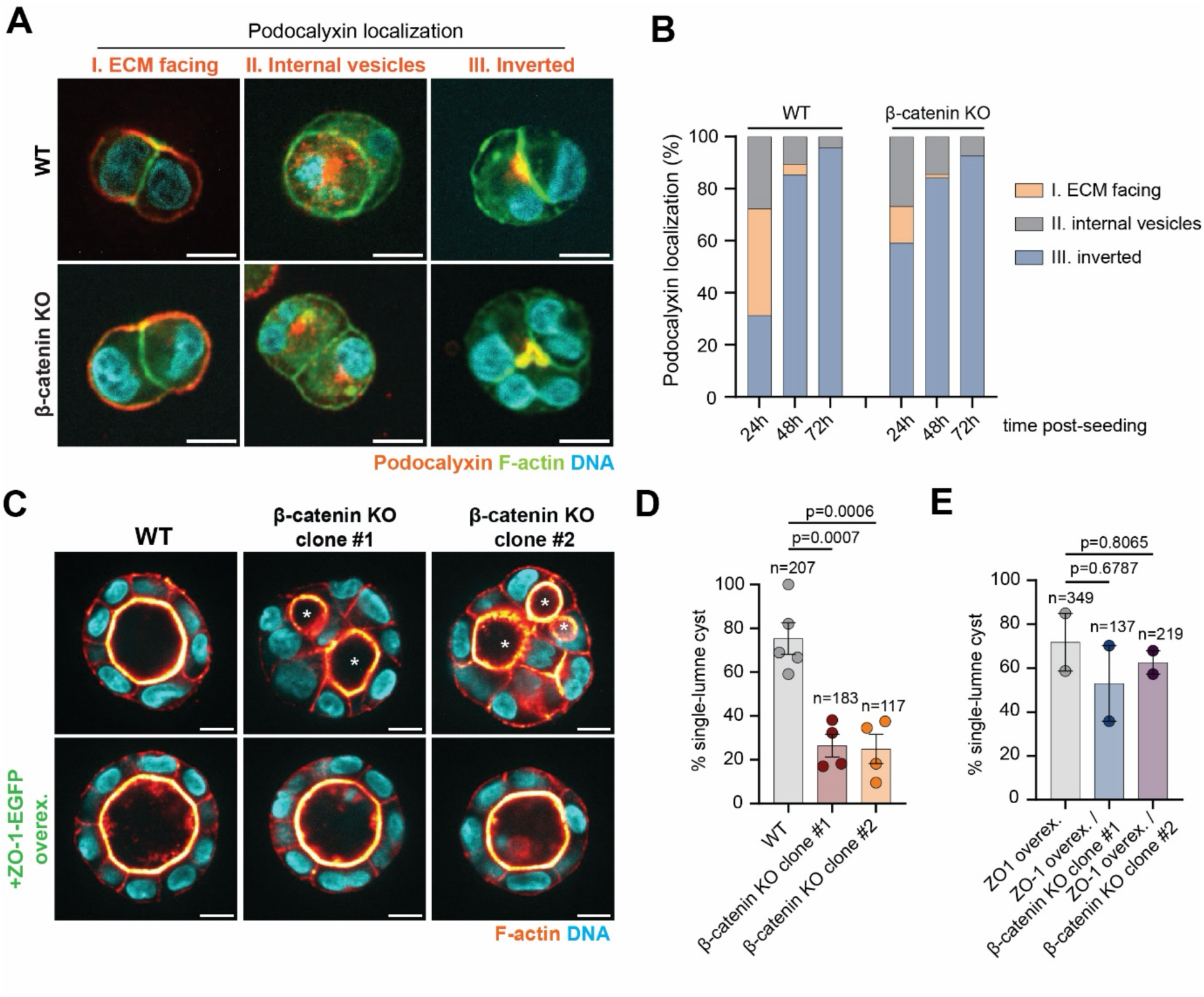
β-catenin-dependent planar epithelial cell division is required for 3D cystogenesis. (**A**) Early stages of single-lumen formation in MDCK 3D matrigel culture. Representative confocal images of podocalyxin (apical membrane marker), F-actin, and DNA depicting three distinct morphology profiles as defined by podocalyxin localization, comparing WT (top) and β-catenin KO (bottom). (**B**) Classification of cyst morphology at 24 h, 48 h, and 72 h, showing similar profiles of β-catenin KO compared to WT during the initial cystogenesis stages. Total cell doublets used for graph plotting at different stages are: 24 h, 83 (WT), 71 (KO), 48 h, 122 (WT), 76 (KO), 72 h, 24 (WT), 54 (KO). (**C**) Requirement of β-catenin for single-lumen formation in matured cyst. Representative confocal images of F-actin and DNA of cysts at day 7 post-seeding, comparing WT with 2 clones of β-catenin KO, generated in parental background (top row) or ZO-1-EGFP/Histone-H2B-mCherry stable expression background (bottom row). (**D-E**) Quantification of single-lumen cysts corresponding to conditions in (C). Sample sizes, and p-values indicated on graphs. To ensure statistical robustness despite the constraint in Matrigel availability, more than 100 cysts per condition were quantified. Total cysts quantified are indicated on the graph. Scale bars, 10 μm (A, C).

Subsequently, we investigated the involvement of β-catenin in the lumenal expansion phase of cystogenesis by assaying the cyst morphology at day 7 in 3D Matrigel culture, at which point the control cyst is considered mature. As shown in **Figure 8C-E**, we found that multiple lumens (>2) are observed for the majority of β-catenin KO cysts, indicative of defects during lumen expansion in knockout cells, consistent with the spindle orientation defects in 2D monolayer described above. Finally, we test whether the overexpression of ZO-1 is also able to rescue mitotic spindle orientation defects in β-catenin KO in 3D. As shown in (**Figure 8C-E and Supplementary Figure 4A**), we assessed the ability of β-catenin KO generated in ZO-1-GFP background (Saw et al., 2022) to generate single-lumen cyst in 3D Matrigel, using 2 clonal populations. We observed that single-lumen formation is significantly restored, approaching WT level, in contrast to β-catenin KO clones without ZO-1-GFP overexpression in which the majority of cysts were multi-lumen. These results thus corroborate the mechanistic framework proposed in the previous sections for β-catenin-dependent spindle orientation.

## Discussion

Analyses in both developmental and cultured cell systems have revealed multiple, partially overlapping mechanisms that regulate the spatial orientation of the mitotic spindle (di Pietro, Echard and Morin, 2016; Donà, Eli and Mapelli, 2022; Lechler and Mapelli, 2021). While the evolutionarily conserved force-generation core of this machinery is primarily the NuMA-LGN-Gαi complex, at the cell cortex a diverse array of molecular assemblies has evolved to serve as adaptors, tailored for specific tissue architecture or another cellular context. In this study, we uncovered a previously uncatalogued function of β-catenin in regulating the mitotic spindle orientation machinery in a symmetrically dividing epithelial cell model. As outlined in **Supplementary Figure 7A-D**, we show that, through its N-terminal domain, β-catenin mediates this function via the recruitment of Afadin to the basolateral cortex of dividing cells. Earlier Afadin has been shown to recruit LGN to the cortex (Carminati et al., 2016) by binding to the TPR domain of LGN. The role of β-catenin delineated here thus represents a molecular upstream event preceding Afadin recruitment. Subsequent to β-catenin-mediated cortical recruitment, LGN-Afadin complex is formed during early mitosis. Since the binding affinity of LGN to Afadin is weaker than LGN-NuMA, Afadin is expected to be displaced by NuMA upon its release from the nucleus following NEBD (Carminati et al., 2016). In our proposed model, β-catenin-mediated Afadin cortical recruitment may help ensure adequate and symmetric cortical distribution of LGN, and subsequently proper in-plane spindle alignment during metaphase. Conversely, in the absence of β-catenin, basolateral recruitment of Afadin is reduced, leading to asymmetric distribution of LGN at the membrane, which affects NuMA stability at the spindle pole and impairs proper spindle orientation.

Interestingly, although β-catenin is required for cortical recruitment of Afadin, we found that in the absence of β-catenin, overexpression of either ZO-1 or Afadin itself is sufficient to rescue spindle orientation defect. From our biochemical analysis to dissect the functional hierarchy (**Supplementary Figure 6A**), we found that Afadin is recruited by ZO-1 in the absence of β-catenin. We thus interpret these results as implicating a minimum threshold level of cortical Afadin for proper spindle orientation. Afadin is a multivalent protein with additional interaction partners at the plasma membrane including nectin (Fujiwara et al., 2015), α-catenin (Gong et al., 2025), and ZO-1 (Ooshio et al., 2010). In β-catenin KO, the overexpressed Afadin could thus be recruited to the cortex via interaction with these partners. Likewise, in β-catenin KO, the overexpressed ZO-1 could serve to recruit Afadin to the cortex in lieu of β-catenin. In this model, in WT or β-catenin KO rescued with ZO-1 or Afadin overexpression (**Supplementary Figure 7C-D**), sufficient Afadin is able to accumulate at the cortex. This enables sufficient recruitment of LGN, resulting in a symmetric distribution of LGN at the cortex. In turn, this allows LGN-NuMA assembly to proceed normally, permitting proper in-plane spindle orientation. On the other hand, in unrescued β-catenin KO, although partial recruitment of Afadin via other partners may occur, the amount of LGN recruited may be insufficient for uniform coverage (**Figure 4C**), hence handicapping the subsequent LGN-NuMA assembly, and causing spindle orientation defect (**Supplementary Figure 7B**).

The existence of the tripartite compensatory network of β-catenin/ZO-1/Afadin suggests that the cortical enrichment of LGN may be a critical system parameter that can be regulated via Afadin-dependent mass action. While our biochemical analysis indicates that β-catenin co-immunoprecipitate with both ZO-1 and Afadin (**Supplementary Figure 6A-C**), such cortical complexes likely contain additional components. In this study, we investigated the role of PAK4, a serine/threonine kinase which has also been shown to directly interact with both Afadin and β-catenin, as well as phosphorylating β-catenin S647 in TCF/LEF signaling pathway (Baskaran et al., 2021; Li et al., 2012). Previously, inhibition of PAK4 kinase activity by PF-3758309 was found to cause aberrant spindle and misaligned chromosomes in oocyte division (He et al., 2019), consistent with the observed chromosome misalignment in MDCK (**Supplementary Figure 2**). However, we found that the spindle orientation was not significantly affected upon PAK4 kinase inhibition, suggesting that the PAK4 kinase activity may be dispensable for proper spindle orientation. As PAK4 may also play a scaffolding role, further study using truncation or mutated variants will be needed to dissect its structural contribution.

The heterohexameric structure of LGN-NuMA complex has been determined recently (Pirovano et al., 2019). The formation of such higher order organization likely requires sufficiently high local concentration of LGN. Given that NuMA itself is known to undergo phase separation (Sun et al., 2021), an intriguing possibility for future study would be whether cortical Afadin-LGN may be present as condensate as well, possibly to promote LGN-NuMA heterohexamerization necessary for anchoring the astral microtubules. Consistent with this, a recent study has shown that β-catenin is capable of forming phase-separated condensates (Monster et al., 2025). Indeed, both ZO-1 and Afadin have also been shown to contain intrinsically disordered region and capable of forming condensates (Beutel et al., 2019; Kuno et al., 2025).

Competitive interaction between NuMA and various cortical scaffolding proteins such as Afadin, Dlg1, E-cadherin, or Inscuteable for the TPR domain of LGN appears to be a common regulatory mechanism that orchestrate distinct steps in mitotic spindle assembly and orientation (Culurgioni et al., 2011; Gloerich et al., 2017; Saadaoui et al., 2014; Yuzawa et al., 2011). In MDCK in particular, E-cadherin has been shown to recruit LGN to regulate mitotic spindle orientation, with the displacement of LGN from E-cadherin by site-specific mutation (758A) causing spindle orientation defect (Gloerich et al., 2017). Indeed, such direct recruitment by E-cadherin likely accounts for partial LGN presence at the cortex in β-catenin KO. We note that the observation from Gloerich et al. study is consistent with the framework of our proposed model in which the critical amount of LGN must be recruited to the early mitotic cortex to ensure symmetric distribution. Here, any loss of cortical LGN recruitment, whether by E-cadherin mutation or β-catenin ablation, could be expected to generate similar outcome that leads to spindle orientation defect. Consistent with our model, the re-expression of junctional proteins ZO-1 and Afadin restore this cortical LGN pool in the absence of β-catenin (**Figure 4C**) to maintain proper spindle orientation. We note that in conventional model of spindle orientation (di Pietro, Echard and Morin, 2016; Kiyomitsu, 2019), LGN is primarily localized at the cortex. However, in this study we observed both the cortical and spindle pole localization of LGN (**Supplementary Figure 3**). Indeed, dual localization of LGN at both the cortex and spindle poles have previously been reported in several studies using MDCK (Zheng et al., 2014; Zheng et al., 2010) or HeLa cells (Fankhaenel et al., 2023; Takayanagi et al., 2019). Such cell-type-dependent differences in LGN distribution may implicate additional previously undocumented roles of LGN and should be further clarified in follow-up studies.

E-cadherin/LGN interaction is structurally analogous to the binding of E-cadherin cytodomain to armadillo catenins, raising a question whether β-catenin could interact with NuMA directly. We first explored this hypothesis computationally by aligning the relevant fragments of β-catenin^135-671^ (Huber and Weis, 2001) or γ-catenin (plakoglobin)^111-676^ (Choi et al., 2009) with NuMA^1861-1928^ (Rizzelli et al., 2020) using ColabFold structure-prediction tool (Mirdita et al., 2022). As shown in a pLDDT plot (**Supplementary Figure 8**), (predicted Local Distance Difference Test, score representing confidence in predicted 3D structure of each amino acid residue within a protein), the prediction for β-catenin^135-671^ reveals a high-confident structured domains spanning from 200-600 residues. However, NuMA^1861-1928^ displays relatively low scores, suggestive of a highly flexible or disordered region, as expected from the short sequence length. Complementing the pLDDT plot, the Predicted Aligned Error (PAE) plot indicates the potential spatial arrangement of residues within the β-catenin^135-671^ and NuMA^1861-1928^, more specifically, the zoomed blue region shows the high confidence of the model in positioning of residue pairs relative to each other in NuMA (Trp55 to Asn65) with β-catenin (Ala391 to Arg565). In contrast, for γ-catenin (plakoglobin)^111-676^ and NuMA^1861-1928^ ColabFold analysis reveals lower confidence. Altogether, this analysis raises the possibility that β-catenin may preferentially interact with NuMA compared to γ-catenin (plakoglobin). Consistent with our computational prediction, β-catenin–NuMA interaction was experimentally validated by co-immunoprecipitation (**Figure 5D**). Notably, Eli et al. recently reported that β-catenin also mediates spindle orientation through the LRP6/Wnt pathway via NuMA/LGN in HeLa cells(Eli et al., 2025). While NuMA does not exhibit a comparable cortical signal in the MDCK context as it does in HeLa cells, likely due to cell-type dependent variation, our findings nonetheless converge on β-catenin as a key regulator of NuMA-dependent spindle orientation, and provide new insight into how it contributes to this process through a distinct, junctional role.

Finally, our findings underscore the wide-ranging, multifaceted nature of β-catenin function, elucidating an essential role beyond its established functions in mediating cell–cell junctions and Wnt-signaling. By serving as an upstream regulator of the Afadin–LGN machinery, β-catenin is indispensable for defining the proper cell division plane in both 2D simple epithelial and 3D cyst models. Specifically, our study reveals that β-catenin governs metaphase spindle orientation through a junctional interplay with ZO-1 and Afadin, maintaining the cortical pool of Afadin required for proper LGN recruitment and spindle alignment, thereby linking cell-cell junction integrity to the mitotic apparatus. While our study primarily focuses on the metaphase spindle orientation, Afadin has also been implicated in division-axis correction at later mitotic stages, including telophase (Lough et al., 2019). Thus, cell-cell junctional proteins continually coordinate spindle and division-plane fidelity across multiple phases of mitosis through distinct yet interconnected mechanisms. Dissecting how these junctional components interact and reorganize over the full course of cell division — ideally through dedicated long-term, high-resolution live imaging — represents an important direction for future work. Another open question concerns the *in vivo* contexts in which β-catenin-dependent AJ–mitosis crosstalk operates. Future studies delineating this pathway in physiological and pathological settings may reveal how its dysregulation contributes to tissue disorganization and tumorigenesis

### Limitations

Several limitations of our study should be noted. First, since mitotic cell division is a dynamic process involving continual changes of intricate sub-cellular machinery, this poses experimental challenge in capturing sufficient number of relatively rare events using high-resolution microscopy. In our study, the spindle orientation was measured at metaphase in fixed cells, to facilitate statistical analysis but at the cost of only capturing only the endpoint of orientation rather than its dynamics. Kinetic differences — such as a prolonged search-and-capture period in rescue cells with partially restored astral microtubules — would not be detected in this analysis and will require future live-cell imaging to address. Second, although the β-catenin KO strategy enable functional dissection of distinct domains of β-catenin, endogenous level expression is challenging to achieve without a knock-in strategy that may hamper the assessment of rescue. For example, while ectopic re-expression of β-catenin in β-catenin KO cells was found to restore spindle orientation, a number of other parameters such as astral microtubule number, spindle volume and length, were partially rescued. This could arise from the rescue level of GFP–β-catenin being modestly sub-endogenous or reflecting the pleiotropic functions of β-catenin, which operates across multiple subcellular pools such as cell–cell junctions, the centrosome, the cytosol and the nucleus and each contributing to distinct cellular processes (Valenta, Hausmann and Basler, 2012). In particular, β-catenin and its destruction-complex partners (APC, Axin, GSK3β) have centrosome-associated roles in microtubule nucleation, centrosome cohesion and separation(Bahmanyar et al., 2008; Fumoto et al., 2009; Hadjihannas, Bruckner and Behrens, 2010) and may therefore influence astral microtubules and spindle geometry independently of the cortical Afadin/LGN pathway that primarily governs orientation. Third, we note that junctional E-cadherin was found to be modestly reduced (∼10%) in β-catenin KO cells. Although complete loss of E-cadherin still permits robust spindle orientation in MDCK cysts(Otani et al., 2019), a partial contribution of altered E-cadherin to our phenotype cannot be entirely excluded. The variability of spindle angles within the knockout population is also consistent with spindle orientation being inherently probabilistic and shaped by multiple, partially redundant junctional inputs, as seen in prior E-cadherin perturbation studies (Gloerich et al., 2017). Fourth, while our co-immunoprecipitation and rescue data support a model in which β-catenin, ZO-1 and Afadin operate as a partially redundant, cooperative scaffold for cortical LGN recruitment rather than as strictly independent pathways, the precise upstream–downstream architecture of this network is not fully resolved. Given that β-catenin loss reduces but does not abolish cortical Afadin, ZO-1 overexpression can restore cortical Afadin in β-catenin KO cells, and all three proteins co-precipitate **(Figures 6C and 7A–B; Supplementary Figure 6A–C**), collectively these suggest that these proteins may be distributed across overlapping rather than entirely separate pools. Dissecting the relative contributions of these scaffolds, and whether they establish distinct sub-cortical sub-pools of Afadin and LGN will require further investigation, ideally with higher-resolution imaging and acute perturbation approaches. Fifth, we note that NuMA is found to be predominantly localised at spindle poles, with any cortical pool below the detection limit of our imaging conditions. While this is consistent with previous studies in MDCK (Zheng et al., 2014; Zhu et al., 2011), this localization contrasts with keratinocytes study (Seldin et al., 2013), and likely reflects cell-type-dependent differences in NuMA distribution. A similar cell-type specificity may explain why we did not observe the monoastral spindle/centrosome-separation defects previously reported in β-catenin-depleted HEK293 cells (Kaplan et al., 2004). Together, these considerations refine the interpretation of our findings: β-catenin acts at the lateral cortex, together with ZO-1 and Afadin, to promote LGN/NuMA-dependent spindle orientation, while its additional functions at the centrosome and other subcellular sites likely account for aspects of the mitotic phenotype that are not restored by junctional rescue alone.

## Materials and Methods

### Cell Culture

MDCK II (Madin-Darby Canine Kidney) cells used were originally characterized by W. James Nelson (Stanford University) and gifted by Chwee Teck Lim (Mechanobiology Institute, NUS). MDCK catenin knockout cell lines (single knockout: β-catenin, γ-catenin and β-/γ-cat double knockout) were generated from parental MDCK II cells(Saw et al., 2022). MDCK II cell line with stable ZO-1-GFP expression was generated by transfecting EGFP–ZO-1 into MDCK cells that stably express mCherry–H2B, a gift from Chin-lin Guo, Academia Sinica, Taiwan (Ouyang et al., 2021) using Lipofectamine LTX (Thermo Fisher Scientific). Stable double-expressing clones were selected by serial dilution under G418 selection (Gibco, 10131). Cells were maintained at 37 °C in a humidified atmosphere with 5% CO₂ and cultured in MEM (Gibco, 11095) supplemented with 10% fetal bovine serum (Hyclone, SH30071.03), 10 mM sodium pyruvate (Gibco, 11360), 100 IU/mL penicillin, and 100 µg/mL streptomycin (Gibco, 15140122).

### Expression vectors and transfection

The fluorescent protein (FP) fusion constructs, γ-catenin-mEmerald (C), was created in the laboratory of Michael W. Davidson, The Florida State University, and available from Addgene repository (#54133). β-catenin insert was cloned from the mEos2 (N)-β-catenin construct, then fused with pmEmerald (C) backbone to obtain β-catenin-mEmerald (C) construct. The mEos2 (N)-β-catenin and pmEmerald (C) backbone are from Michael W. Davidson. The subsequent truncation constructs of γ-cateninΔN1-134-mEmerald (C), γ-cateninΔC671-781-mEmerald (C), β-cateninΔN1-124-mEmerald (C), β-cateninΔC669-745-mEmerald (C) were generated from the full length following the amino acids information from previous publication(Aberle et al., 1996). β-catenin point mutation constructs, Y142E/Y142F and β-catenin Δα-catenin binding (aa: 118-169) are synthesized by Epoch Life Science, Inc. Afadin-mcherry is gift from Marina Mapelli lab. Afadin dominant negative form (S216A/S1090A) was introduced by point mutations based on the Afadin-mcherry vector and synthesized by GentleGen. EB1-GFP (#39299) and ZO-1-EGFP (#30313) are from Addgene Davidson Collection. Unless indicated otherwise, plasmids were transfected by electroporation (Neon transfection system, Life Technologies) according to manufacturer’s protocols. The β-catenin full-length (FL)-mNeonGreen construct was generated in-house by subcloning the β-catenin-FL coding sequence from a β-catenin-FL-mEmerald plasmid into the pLV-hTERT-IRES-hygro lentiviral backbone (a gift from Rong Li, Mechanobiology Institute, National University of Singapore), with mNeonGreen replacing mEmerald as the fluorescent tag. Lentiviral particles were packaged according to the manufacturer’s protocol using the Lenti-X™ Packaging Single Shots system (Takara Bio, #631275) and used to transduce target cells. Three days post-transduction, single-cell sorting was performed on a Sony SH800 cell sorter using the FITC channel to isolate mNeonGreen-positive clones, which were subsequently expanded in culture until ready for use.

### Generation of armadillo catenin knockout MDCK II cell lines

Bi-allelic knock-out of β-catenin, and γ-catenin were generated by CRISPR/Cas9-mediated genomic editing using dual sgRNAs to induce exon skipping. The dual sgRNA were designed with CHOPCHOP web tool (https://chopchop.cbu.uib.no) and synthesized by Integrated DNA Technologies (IDT). Each guide sgRNA sequence is designed to flank the early exon of respective *Canis familiaris* genes, with the following sequences.

The pairs sgRNAs target to β-catenin are sgRNA-A2-5’-GCCAAACGCTGGACATTAGTAGG-3’ and sgRNA-B2-5’-CACAACCGAATCGTAATCAGAGG-3’. The pair sgRNAs target to γ-catenin are sgRNA-A3-5’-GTCCGATGCTTGGGGTGTAG-3’ and sgRNA-B3-5’-ACCTGACGTGCAACAACAGC-3’. The oligo sequences were ligated into the pSpCas9 (BB)-2A-Puro (PX459) V2.0 backbone plasmid (Feng Zhang laboratory, Addgene #62988), which contains Cas9 gene with BbsI cloning site under the control of U6 promoter. The hU6-F primer (5′-GAGGGCCTATTTCCCATGATT-3′) was used for sequencing to confirm the presence of sgRNA oligos after cloning (Axil Scientific Sequencing). MDCK II cells (5 × 10^5^ cells per transfection) was electroporated with 5 µg of plasmids containing sgRNA-A and sgRNA-B, by utilizing a Neon™ Transfection System (100 μL tips, Thermo Fisher #MPK10096) with pulse rate of 1650 V, 20 ms pulse width, and pulse number of 1. The transfected cells were grown for 48-hour before selection using 2 µg/mL puromycin (Sigma-Aldrich #P8833). Following puromycin selection, cells were suspended and cultured in 96 wells with single cell per well. Genomic DNA was then extracted for each well and PCR was performed to detect deletion band and non-deletion band, using primers flanking the CRISPR cut sites and primers in the excised region, respectively. Bi-allelic deletion single-cell clones are selected from wells which exhibit deletion bands and non-deletion bands. Upon cell growth, DNA extraction, and PCR was performed to confirm biallelic deletion. To generate the β-/γ-catenin double knockout, the Alt-R CRISPR-Cas9 System editing was performed on the β-catenin knockout cells. The AltR CRISPR-Cas9 crRNA was mixed with tracerRNA 550 in equimolar concentration of 200 µM. Upon addition of the Alt-R Cas9 enzyme, the mixture was used for transfection by Neon electroporation system as described above. Colored tracerRNA was co-transfected to enable selection of positive colonies by FACS sorting. Clones were picked from 96-well plates, amplified and validated for bi-allelic deletion by immunofluorescence microscopy and western blotting. The γ-catenin crRNA sequences used are as crRNA2-/AltR1/rCrU rArCrA rCrCrC rCrArA rGrCrA rUrCrG rGrArC rGrUrU rUrUrA rGrArG rCrUrA rUrGrC rU/AltR2/, crRNA3-/AltR1/rGrC rUrGrU rUrGrU rUrGrC rArCrG rUrCrA rGrGrU rGrUrU rUrUrA rGrArG rCrUrA rUrGrC rU/AltR2/, crRNA6-/AltR1/rGrC rArCrA rUrCrA rCrUrC rGrArG rUrArC rArUrG rGrUrU rUrUrA rGrArG rCrUrA rUrGrC rU/AltR2/.

### RNA interference

For siRNA-mediated knockdown of Afadin in MDCK II cells, Dicer Substrate siRNA (DsiRNA) pairs designed using IDT online tool was applied to target the Afadin CDS sequence (Gene ID: 431695) in the *C. familiaris* genome. 10 μM stock solution was prepared with Nuclease-Free water (UltraPure^TM^ Distilled Water, Invitrogen #10977-015) and denatured at 94 °C for 2 min, aliquoted, and stored at −80 °C. Sequences used for targeting Afadin are dsiRNA1 rGrArU rGrArC rArUrU rCrCrA rArArU rArUrA rArArC rArGT A and dsiRNA2 rArArG rArUrA rArArC rUrUrG rGrArA rUrCrU rArUrG rUrCA A. Both siRNAs are stored in −20 °C according to the manufacturer’s protocols. The siRNA pairs were mixed (to a final 20nM working concentration) and co-transfected with TYE563 transfection control DsiRNA using Lipofectamine™ 3000 (Invitrogen MAN0009872) to visualize the transfection efficiency. The immunostaining for spindle observation was carried out after a 48-hour post transfection.

### Antibodies and reagents

Primary antibodies, Rabbit β-catenin (ab2365, 1:200), rabbit α-catenin (ab51032, 1:400), mouse α-tubulin (ab7291, 1:200), rabbit α-tubulin (ab18251, 1:200), rabbit γ-tubulin (ab179503, 1:200) were purchased from Abcam. Mouse γ-catenin/plakoglobin (138500, 1:400) and rabbit ZO-1 (61-7300, 1:400) were purchased from Invitrogen. Mouse γ-tubulin (T6557, 1:200) was from Sigma. Rabbit LGN (ABT-174, 1:200) was from Millipore. Mouse Afadin/AF-6 (MAB78291, 1:200) was from Biotechne. Rabbit β-catenin pSer33/37/Thr41 (9561, 1:200) was from Cell Signaling. Mouse podocalyxin/gp135 (3F2/D8-s, 1:200) was requested by Developmental Studies Hybridoma Bank (DSHB). Mouse E-cadherin (3B8, 1:200) was a gift from Dr. Alpha S. Yap (The University of Queensland). Rabbit NuMA (1:200) was a gift from Dr. Duane A. Compton (Dartmouth Geisel School of Medicine). Secondary antibodies, donkey anti-rabbit IgG Alexa Fluor^TM^ 568 (A10042, 1:200), donkey anti-mouse IgG Alexa Fluor^TM^ 647 (A31571, 1:200) were purchased from Invitrogen. Phalloidin-ATTO488 (49409, 1:200) was from Bioreagent. Nucleus dye Hoechst 33342 (62249, 5ug/ml) was from ThermoFischer Scientific.

### Immunofluorescence microscopy

For immunostaining, 2D monolayer of cells were fixed with freshly prepared 4% formaldehyde (Electron Microscopy Sciences) in PBS for 15 min and then permeabilized with 0.1% Triton X-100 in PBS for 10 min at room temperature. 3D Matrigel cultured cysts were fixed under the same condition, but with the fixation duration extended to 30 min, and with permeabilization performed using 0.2% Triton X-100 in PBS for 30 min at room temperature. Followed by blocking with 4% BSA (Sigma-Aldrich #A7906-100g) for 1 hour at room temperature. Fixed cells were first incubated with primary antibody overnight at 4°C, followed by secondary antibody incubation with or without staining by DAPI, Hoechst, or Phalloidin for 1 hour at room temperature. Between each step, cells were washed thrice with PBS for 5 min at room temperature each on a rocker.

### Confocal Microscopy

Spinning disk confocal/Super-resolution imaging was performed on a Nikon Eclipse Ti-E inverted microscope, coupled with confocal scanner unit CSU-W1 (Yokogawa), super-resolution module Live-SR (Gataca Systems), Prime95B scientific CMOS camera (Photometrics) and controlled by MetaMorph software (Molecular Devices). Live cells were imaged with 60X/1.2 CFI Plan Apochromat VC water immersion objective at 37 °C 5% CO_2_, and fixed cells with 100X/1.45 CFI Plan Apochromat λ oil immersion objective. The dichroic and emitters on W1-Spinning Disk are as follow: Pinhole size: 50um, Dichroic: 405/488/561/ 640, Emitter for Ex 405nm: 445/45, Emitter for Ex 488nm: 525/40, Emitter for Ex 561nm: 617/73, Emitter for Ex 642nm: 692/40.

### 3D matrigel culture

To keep cells in highly proliferative stage, MDCK II cells were replated in T25 flask and let grow to around 60-70% confluence before tryptic digestion. Cells were seeded as single-cell suspension in 2% Matrigel (10^4^ cells) on 100% Matrigel (ThermoFisher #A1213302) coated glass bottom dish (Iwaki, diameter 12/27mm). Culture medium was refreshed every two days before fixation. For the pre-apical podocalyxin transcytosis assay, 3D cysts were fixed at 24-hour, 48-hour, and 72-hour to observe podocalyxin localization. For matured cysts observation, cysts were fixed at day 7 post seeding. The percentage of single lumen cysts was calculated by dividing the multiple lumen cysts numbers by the total number of cysts. MDCK II podocalyxin transcytosis assay was performed on fixed sample post seeding from 24/48/72-hour to characterize the early AMIS formation processes.

### Mitotic spindle profile and orientation angle analysis

To analyse mitotic spindle orientation, cells were plated on Iwaki glass-bottom dishes, let grow to reach confluence, and then fixed and stained for γ-tubulin to visualize spindle poles, α-tubulin for spindle structure and nuclear. Metaphase mitotic spindle profile is analysed using the Spindle3D plugin in ImageJ.

### Quantitative analysis of protein spatial distribution

To quantify the cortex-associated intensity of LGN or Afadin cell boundary was segmented using the p120-catenin immunostaining channel to define the region of interest, background signal is measured by the blank region on the images, all intensity measured is subtracted from the background. The analysis of tight-junction integrity was performed by comparing the length of tight junctions visualized by ZO-1 immunostaining with p120-catenin immunostaining of adherens junction. To quantify NuMA intensity at the mitotic spindle, a binary mask corresponding to the mitotic spindle region was generated by segmentation of the α-tubulin channel. The relative NuMA signal was calculated as the ratio of NuMA to α-tubulin fluorescence intensity within this mask.

### Microtubule dynamics analysis

To monitor microtubule dynamics, microtubule plus-ends were labelled by transfecting cells with an EB1-GFP construct using Lipofectamine 3000 (Cat. #L3000008, Thermo Fisher Scientific). Transfected cells were replated at 24 h post-transfection and imaged at 48 h. Live EB1 comets are imaged using BC43 Benchtop confocal under 60x lens for 100ms intervals over 30s. Plus-end tracking velocities were analysed using the TrackMate plugin in ImageJ (Dixit et al., 2009) with the LoG detector in segmentation comets spots (estimated object diameter is 0.6 micron, quality threshold is 6) and Advanced Kalman Tracker in frame-to-frame linking (initial search radius is 0.2 micron, search radius is 0.8 micron and max frame gap is 3).

### Immunoblotting and Co-immunoprecipitation Assay

For western blot analysis, cells were collected and lysed in Pierce RIPA buffer (Cat. #89900, Thermo Fisher Scientific) supplemented with proteinase inhibitor (Cat. #5871, Cell Signaling Technology). Protein concentrations in the lysate were determined using the Pierce BCA Protein Assay Kit (Cat. #A55864, Thermo Fisher Scientific). 15 μg of whole cell lysates were resolved by SDS-electrophoresis (Bio-Rad Mini-Protean Tetra Cell, Cat. #1658029 and 4-20% Cat. #4561094) and transferred to 0.2 μm PVDF membrane (Bio-Rad Cat. #1704156) with Trans-Blot Turbo Transfer System (Bio-Rad Cat. #1704150). Blocking was performed in Intercept (TBS) Blocking Buffer (Cat. #92760001). Primary antibody was diluted in 50% Blocking buffer (in TBST) and incubation was performed at 4 °C for 16 h. Secondary antibody was incubated for 1h at room temperature before imaging.

For ZO-1 and Afadin pull-down assays, MDCK rescued expression full-length β-catenin cells were harvested at 80–90% confluence and lysed in Pierce RIPA Buffer (Cat. #89900, Thermo Fisher Scientific) supplemented with proteinase inhibitor (Cat. #5871, Cell Signaling Technology). Protein concentrations in the lysate were determined using the Pierce BCA Protein Assay Kit (Cat. #A55864, Thermo Fisher Scientific). For each pull-down, 500 μg of whole-cell lysate was incubated with either ChromoTek mNeonGreen-Trap Agarose (Cat. #nta-20) or GFP-Trap Agarose (Cat. #gta) according to the manufacturer’s instructions. To stabilize intracellular interaction complexes, cells were treated with the cross-linker DTBP (dimethyl 3,3′-dithiobispropionimidate; Cat. #D2388, Merck) for 30 min prior to lysis, followed by quenching with 100 mM Tris-HCl (pH 8.0) for 10 min on ice before harvesting.

For the NuMA pull-down assay, MDCK cells were synchronized at the G2/M boundary by treatment with 9 μM RO-3306 (Enzo Life Sciences) for 16 h. Cells were then released for 30 min, treated with 10 μM MG132 (Bio-Techne) for 1 h, and harvested for downstream analysis.

### Statistical analysis

All experiments were repeated at least three times, unless otherwise indicated. Each experiment contained all the wild-type and knockout under the same condition. Results are expressed as mean values ± S.E.M. Individual data points were plotted with *P*-values written on top of the graph. Data from individual groups was first analysed for normal and lognormal distribution. If all groups were normally distributed, a two-tailed t-test was used for 2 groups, and an ordinary one-way analysis of variance (ANOVA), followed by the Tukey’s post hoc test was used for more than 2 groups. If all groups were log-normal, the data was transformed into a normal distribution by taking the log and then analysed. If all groups were neither normal nor log-normal, the Mann-Whitney test was used for 2 groups, and Kruskal-Wallis test, followed by Dunn’s post hoc test was used for more than 2 groups. Statistical analysis was performed using GraphPad Prism version 10.3.1 (GraphPad Software, San Diego, CA, USA).

## Data and Code Availability

Data that supports the plots and figures within this paper as well as analysis codes are available from the corresponding author upon reasonable request.

## Acknowledgements

We acknowledge funding support from Ministry of Education Singapore Academic Research Fund Tier 2 (MOE-T2-EP3-0124-0012, to P.K.), Ministry of Education Singapore Academic Research Fund Tier 3 (MOE-T3-2020-01, to P.K.), MBI intramural funding (P.K.), and National Research Foundation Singapore (NRF-MSG-2023-0001, to P.K.). We thank Jau-Yi Wu (Academia Sinica, Taiwan) for contributing to ZO-1 stable cell line generation. Y.M. is supported by Mechanobiology Institute (MBI) Graduate Scholarship. S.S.C. is partially supported by Final Year Project funding from Department of Biomedical Engineering, National University of Singapore. We thank the microscopy, IT, high-throughput molecular genetics, and wet lab core facilities at MBI for helpful discussion and infrastructure support.

## Authors contribution

Y.M. and P.K. designed the study and wrote the manuscript. Y.M. and P.R. performed the experiments. S.S.C., C.S.L.L., and K.H.L. generated the cell lines used in this study.

## Supplementary Information

**Supplementary Figure 1.**
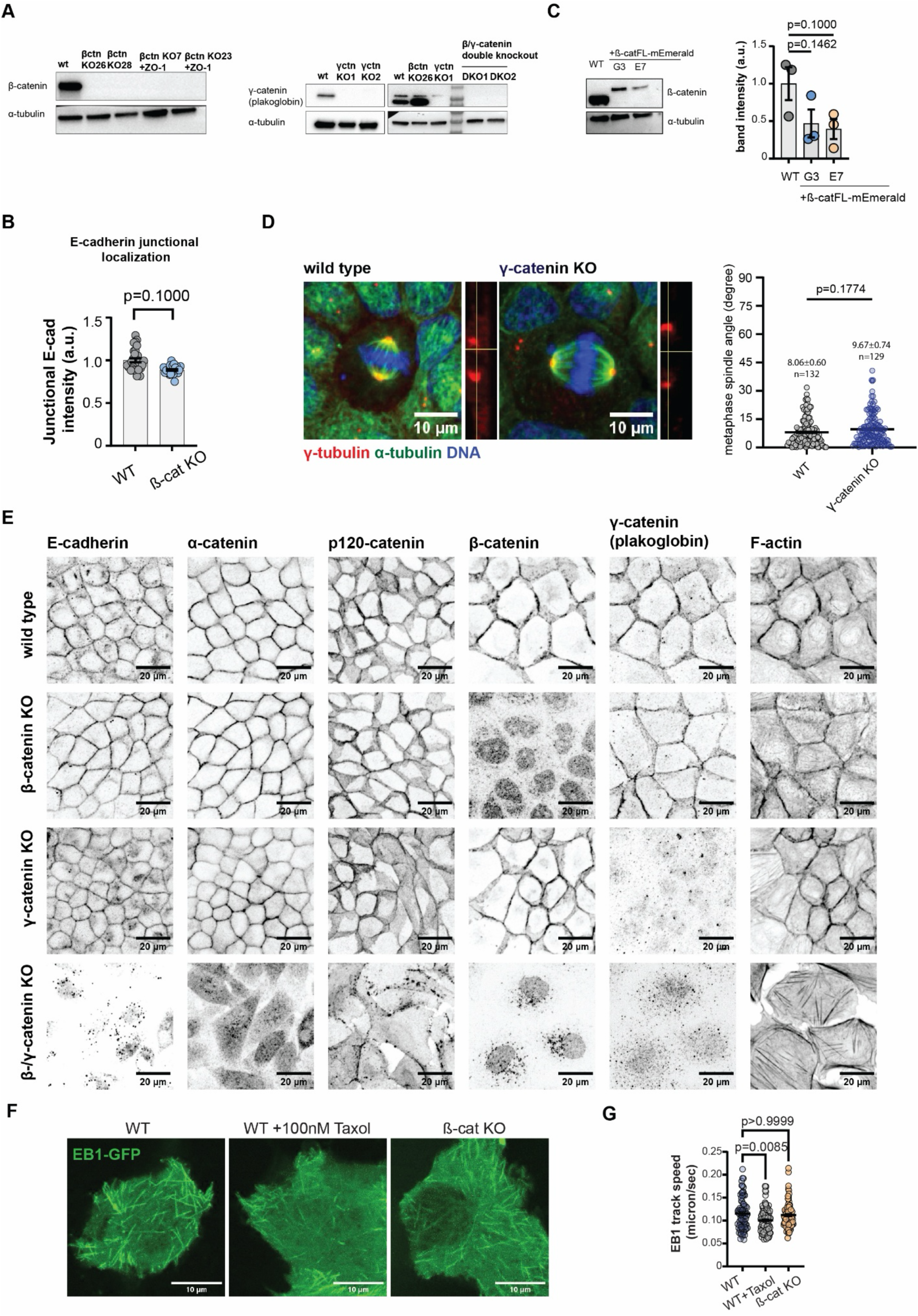
CRISPR/Cas9-mediated ablation of β-catenin and γ-catenin (plakoglobin) (**A**) Validation of CRISPR/Cas9-mediated knockout. (Left) Immunoblot of β-catenin and α-tubulin loading control in MDCK II cells comparing WT, 2 clones of β-catenin biallelic KO generated in parental MDCK II background, and 2 clones of β-catenin biallelic KO generated in ZO-1-EGFP/ Histone H2B-mCherry stable expression background. (Center and Right) Immunoblot of γ-catenin (plakoglobin) and α-tubulin loading control in MDCK II cells comparing WT and 2 clones of γ-catenin biallelic KO (Center) and WT with the clones of β-catenin KO or γ-catenin KO chosen for further use, as well as two double-KO clones (Right). (**B**) Quantification of E-cadherin junctional intensity by immunostaining of WT and β-catenin KO cells probing for endogenous E-cadherin. (**C**) Expression level of β-catenin in rescued clonal cell lines (G3 and E7) compared to WT endogenous levels. Representative immunoblots and quantification of relative band intensity shown. (**D**) Representative confocal images (left) of dividing cells comparing WT and γ-catenin KO, with right inserts corresponding to side view projection in γ-tubulin channels. (Right) Quantification of metaphase spindle orientation angle. Sample sizes, mean ± s.e.m., and p-value indicated on graph. (**E**) Immunofluorescence micrographs of cell-cell junction markers comparing WT and KO cells. (**F**) Representative live-cell images from **Supplementary Movies 1-3** comparing microtubule dynamics as reported by end-binding protein EB1-GFP in WT, WT treated with taxol to perturb microtubule dynamics, and β-catenin KO cells. (**G**) Comparison of EB-1 tracking speed in WT, WT Taxol treated and β-catenin KO cells. Scale bars: 10 μm (D, F), 20 μm (E).

**Supplementary Figure 2.**
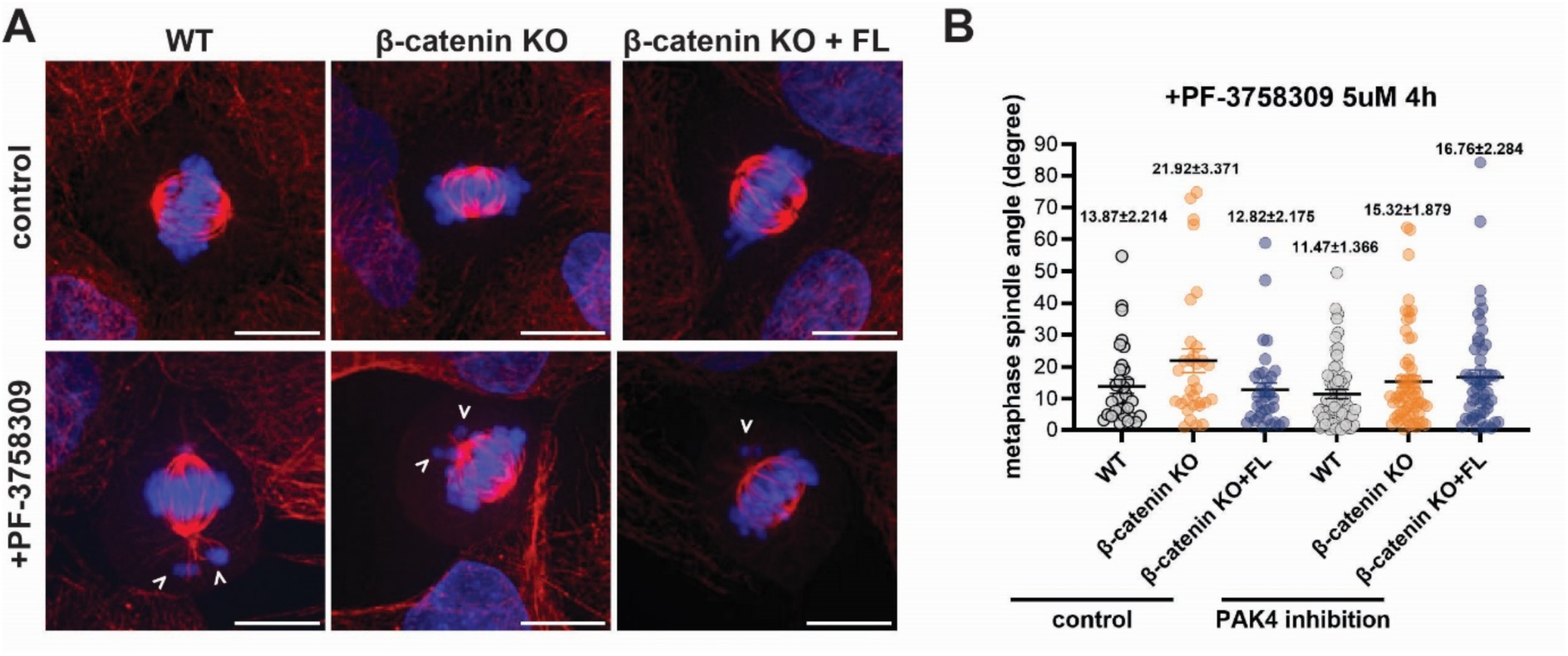
PAK4 inhibition affects chromosome capture but not spindle orientation defects. (**A**) Confocal images of dividing cells comparing WT, β-catenin KO, and KO cells rescued by β-catenin-mEmerald (FL) re-expression, upon treatment by PAK4 inhibitor of β -catenin nuclear shuttling (bottom row, PF-3758309). Arrowheads indicated lagging chromosomes. (**B**) Quantification of metaphase spindle orientation corresponding to conditions in (A). Scale bars: 10 μm (A).

**Supplementary Figure 3.**
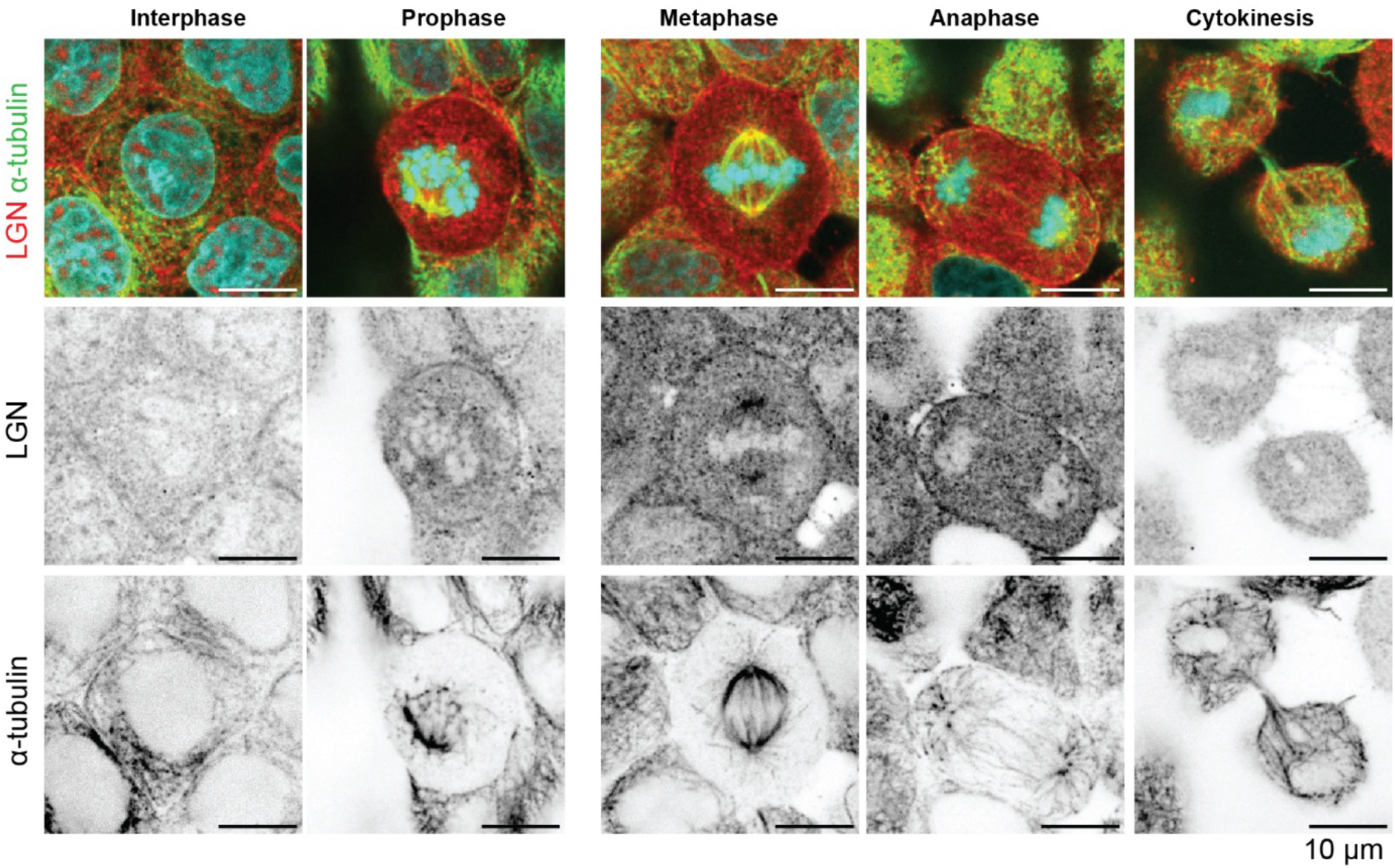
Localization of LGN during cell cycle in MDCK. Representative confocal immunofluorescence micrographs of MDCK II cells across distinct cell cycle stages (interphase, prophase, metaphase, anaphase, and cytokinesis), co-stained for endogenous LGN (red) and α-tubulin (green); DNA was counterstained with DAPI (cyan). Single confocal sections corresponding to the middle plane of each target cell are shown. The top row shows merged channels, while the middle and bottom rows display the individual LGN and α-tubulin channels, respectively, in inverted grayscale to better visualize signal distribution. During interphase, LGN is diffusely distributed throughout the cytosol. Upon mitotic entry, LGN undergoes dynamic redistribution, progressively relocalizing from the cytosol to both the cell cortex and the mitotic spindle. Notably, LGN exhibits dual localization at the cortex and the spindle apparatus during prophase, metaphase, and anaphase, before returning to a predominantly cytosolic pool at cytokinesis. Scale bars, 10 μm.

**Supplementary Figure 4.**
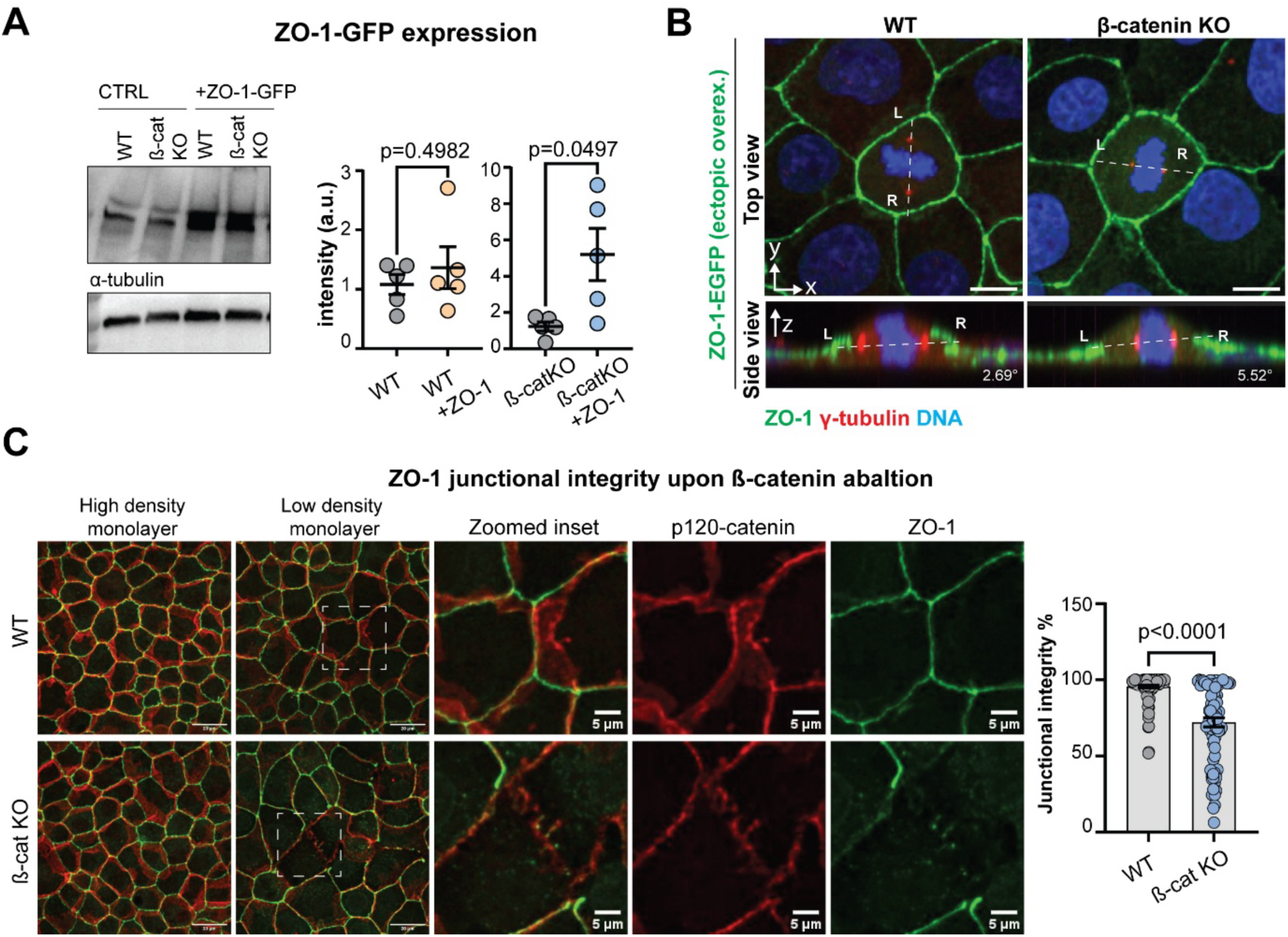
ZO-1 overexpression rescue spindle orientation defects in the absence of β-catenin. (**A**) Representative confocal images of dividing cells with ZO-1-EGFP overexpressed in MDCK WT or β-catenin KO (Fig. 6B main text), (top row). Sideview panels (bottom row) correspond to projection along the dotted lines in top panel, with spindle axis and spindle angle relative to the basal plane indicated. Scale bar, 10 μm. (**B**) Quantification of ZO-1 expression levels between WT and β -catenin KO generated in parental or ZO-1-EGFP stable expression background. (**C**) ZO-1 staining show junctional gaps in the absence of β-catenin. Scale bars: (A), 20 μm and 5 μm (zoomed inset) (C).

**Supplementary Figure 5.**
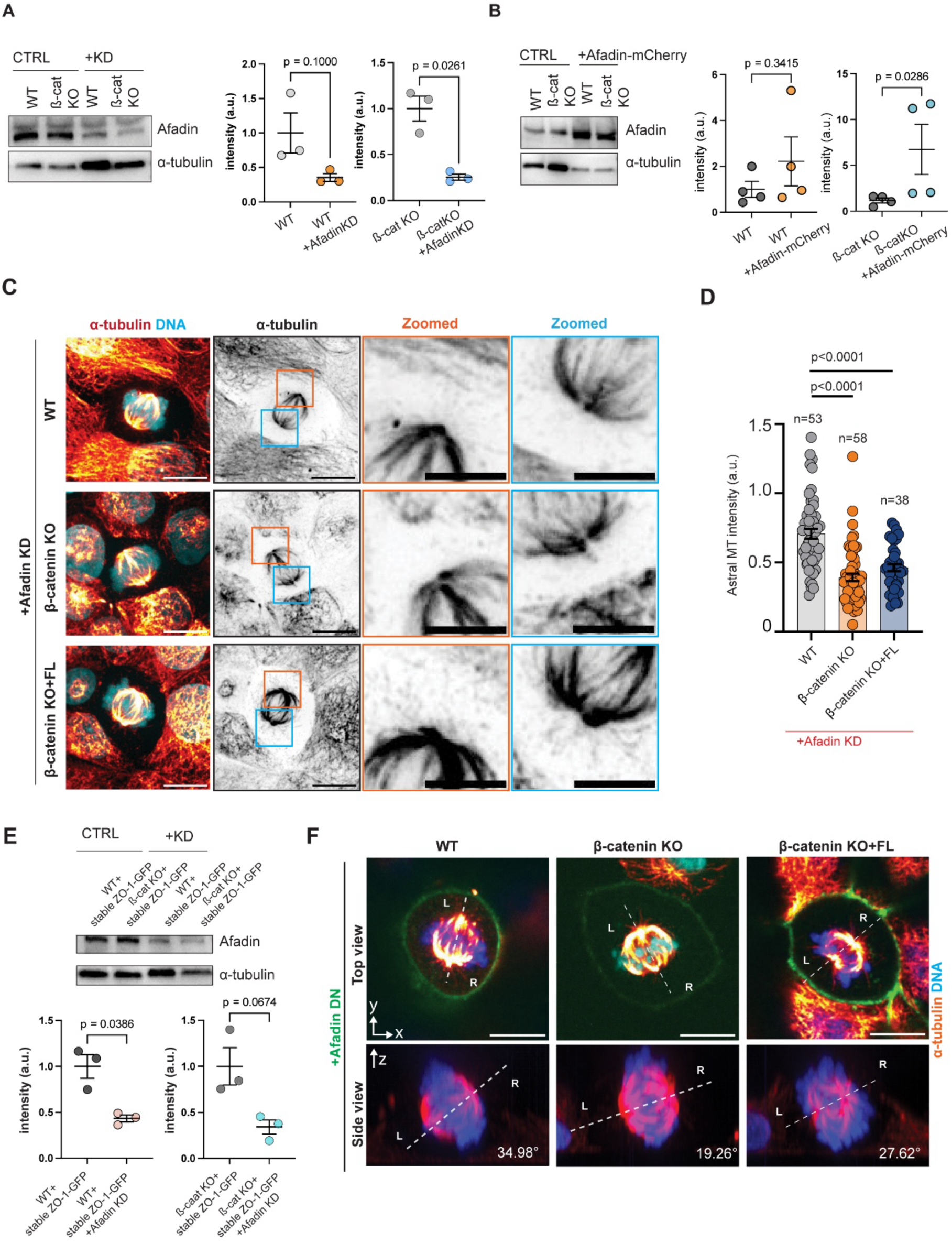
Afadin cortical localization regulates astral microtubules and spindle orientation. (**A**) Depletion of Afadin by dsiRNA. Immunoblots and quantification of Afadin expression levels comparing control with knock-down in WT and β-catenin KO. (**B**) Immunoblot and quantification of Afadin expression levels comparing endogenous (CTRL) with ectopic expression of Afadin-mCherry in WT and β-catenin KO. (**C**) Representative confocal fluorescence micrographs of microtubules and chromosomes in metaphase cells upon Afadin knock down, comparing WT, with β-catenin KO and rescued cells. (Left) Maximum intensity projection 2-color composite (tubulin, DNA), (inner left) α-tubulin channel, (inner right and rightmost) zoomed views corresponding to spindle poles denoted by orange or blue boxes in inner left column. (**D**) Quantification of astral microtubule density. Sample sizes and p-values indicated on graph. (**E**) Depletion of Afadin by dsiRNA in ZO-1-GFP overexpressing background. Immunoblot and quantification of Afadin expression levels, comparing control ZO-1-GFP overexpressing WT and β-catenin KO with knock-down. (**F**) Representative confocal fluorescence micrographs of microtubules and chromosomes of dividing cells, transfected with GFP-tagged dominant negative (DN) Afadin mutation which led to spindle misorientation (Fig. 6E). Sideview panels (bottom row) correspond to projection along the dotted lines in top panel, with spindle axis and spindle angle relative to the basal plane indicated. Scale bar, 10 μm (C,F).

**Supplementary Figure 6.**
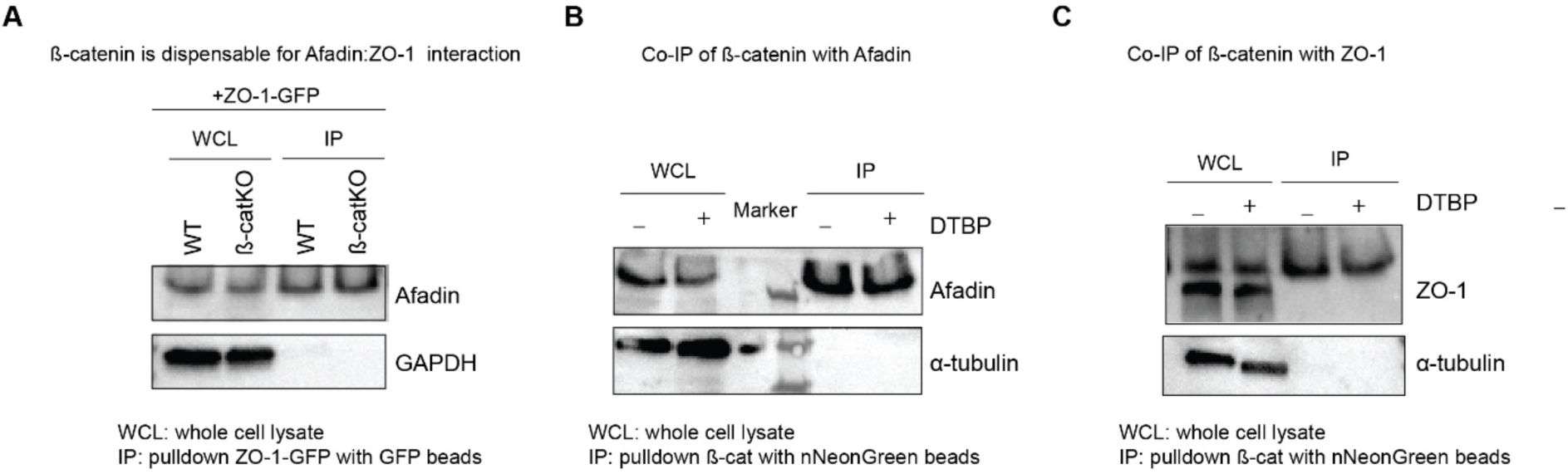
Molecular interaction between Afadin, ZO-1, and β-catenin. (**A**) Immunoblot showing co-immunoprecipitation of ZO-1-GFP with Afadin in both control and β-catenin KO. (**B**) Co-immunoprecipitation of β-catenin-mNeonGreen expressed in β-catenin KO cells with endogenous Afadin under control and DTBP cross-linking. (**C**) Co-immunoprecipitation of β-catenin-mNeonGreen expressed in β-catenin KO cells with endogenous ZO-1 under control and DTBP cross-linking. The experiment was repeated three times.

**Supplementary Figure 7.**
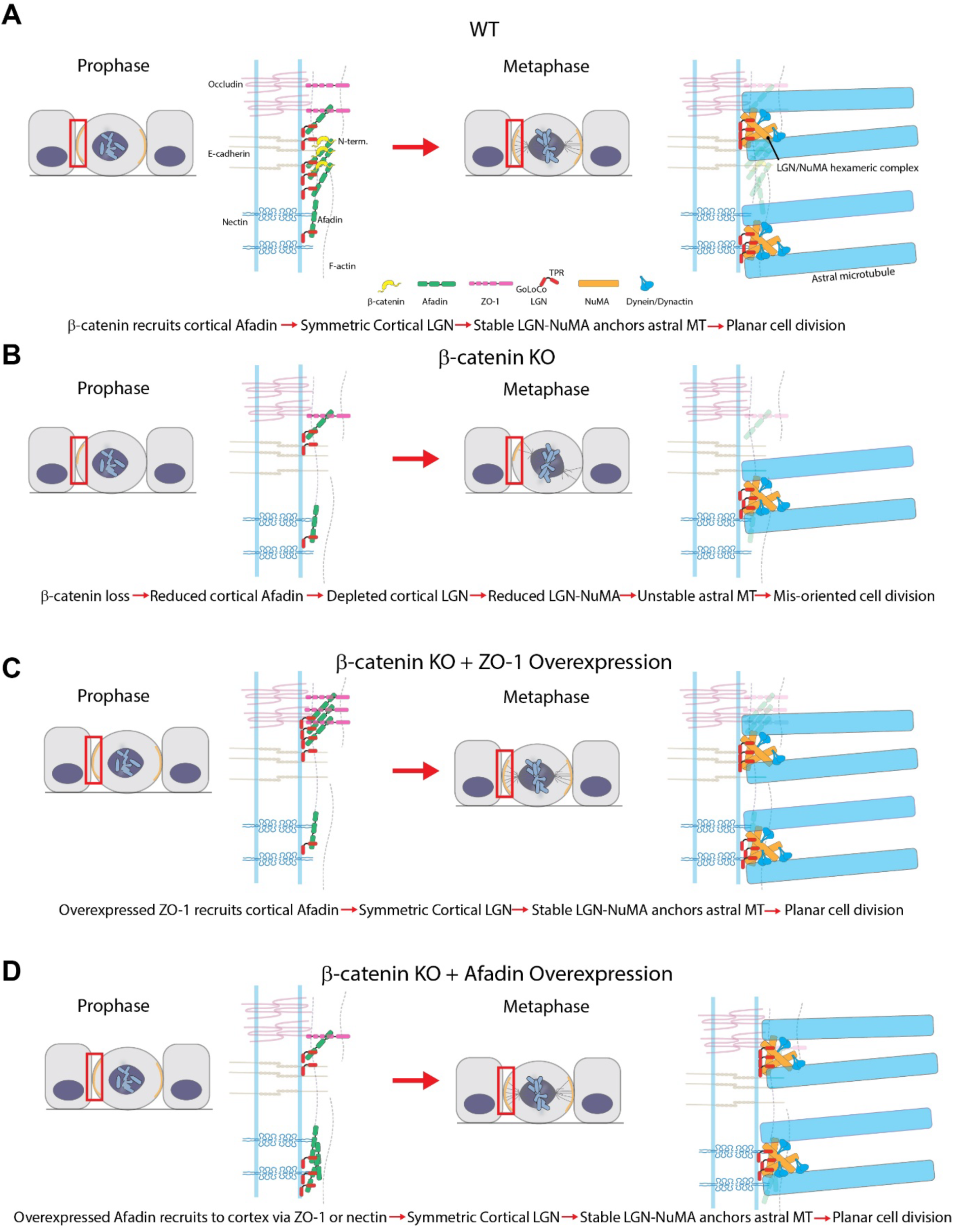
Working model for β-catenin-mediated regulation of symmetric epithelial cell division. **(A)** During early mitosis in WT cells (left), β-catenin (yellow) serves as a major cortical factor that recruits Afadin (green), via its N-terminal domain, which in turn recruits LGN (red). Subsequently, following NEBD, NuMA is released from the nucleus and displaced Afadin from LGN-TPR binding, leading to the formation of heterohexameric LGN-NuMA complex to anchor astral microtubules at the cortex, ensuring planar spindle alignment. (**B**) Upon the loss of β-catenin, depletion of cortical Afadin is associated with reduced LGN cortical accumulation, subsequently leading to deficiency in LGN-NuMA complex at the cortex and hence poor anchorage of astral microtubules. (**C**) Overexpression of ZO-1 in β-catenin KO background is capable of rescuing cortical Afadin due to direct ZO-1/Afadin interaction, leading to adequate LGN. Subsequently, LGN-NuMA complex formation proceeds normally and permits planar cell division. (**D**) Overexpression of Afadin is also capable of compensating for β-catenin KO, since excess Afadin could be recruited by Nectin or ZO-1. Following Afadin cortical accumulation during early mitosis, cortical LGN recruits NuMA after NEBD and permits proper spindle orientation.

**Supplementary Figure 8.**
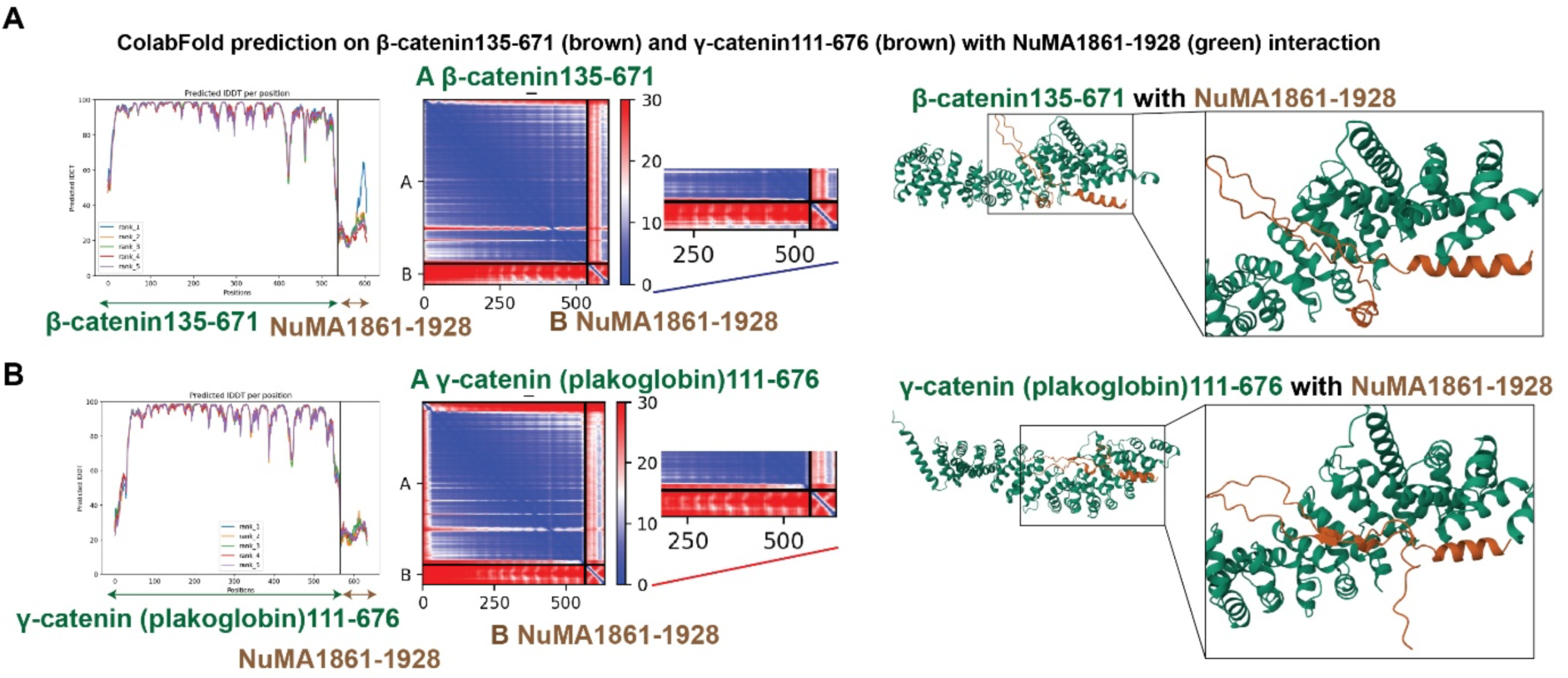
Potential Interaction between β -catenin and NuMA. (**A**) ColabFold prediction of the interaction between β-catenin^135-671^ (brown) and (**B**) γ-catenin^111-676^ (brown) with NuMA^1861-1928^ (green) interaction. Here we evaluate whether direct interaction between NuMA and β-catenin but not γ-catenin may also participate in the regulation of spindle orientation in MDCK II cells. The corresponding E-cadherin binding region of both β-catenin and γ-catenin were aligned with NuMA. The Predicted Aligned Error (PAE) plot (middle panel) indicates the potential spatial arrangement of residues within the β-catenin^135-671^ and NuMA^1861-1928^ in the zoomed blue region shows the high confidence of the model in positioning of residue pairs relative to each other in NuMA (Trp55 to Asn65) with β-catenin (Ala391 to Arg565). There is lower confidence (no blue highlights) in the interaction between γ-catenin (plakoglobin)^111-676^ and NuMA^1861-1928^. Altogether, this analysis raises the possibility of interactions between β-catenin and NuMA.

### Supplementary Videos

**Supplementary Video 1**

EB1 comets in MDCK WT cells. Live imaging of EB1-GFP shows numerous fast-moving comets tracking growing microtubule plus-ends, representing baseline microtubule dynamics (mean tracking speed = 0.14 ± 0.004 μm/sec, n = 78 cells). Related to **Supplementary Figure 1F-G**. Scale bar, 10 μm.

**Supplementary Video 2**

EB1 comets in MDCK WT cells with Taxol treatment. Acute treatment with 100 nM Taxol reduces both the number and tracking speed of EB1-GFP comets, reflecting suppressed microtubule dynamics (mean tracking speed = 0.10 ± 0.003 μm/sec, n = 104 cells). Related to **Supplementary Figure 1F-G**. Scale bar, 10 μm.

**Supplementary Video 3**

EB1 comets in MDCK β-catenin KO cells. EB1-GFP comet density and tracking speed are similar to WT control, indicating that β-catenin loss does not detectably alter microtubule plus-end growth dynamics (mean tracking speed = 0.13 ± 0.003 μm/sec, n = 77 cells). Related to **Supplementary Figure 1F-G**. Scale bar, 10 μm.

## Reference

Aberle, H., H. Schwartz, H. Hoschuetzky, and R. Kemler. 1996. Single amino acid substitutions in proteins of the armadillo gene family abolish their binding to alpha-catenin. The Journal of biological chemistry. 271:1520–1526.

Anjur-Dietrich, M.I., V. Gomez Hererra, R. Farhadifar, H. Wu, H. Merta, S. Bahmanyar, M.J. Shelley, and D.J. Needleman. 2024. Mechanics of spindle orientation in human mitotic cells is determined by pulling forces on astral microtubules and clustering of cortical dynein. Dev Cell. 59:2429–2442 e2424.

Baer, M.M., H. Chanut-Delalande, and M. Affolter. 2009. Cellular and molecular mechanisms underlying the formation of biological tubes. Curr Top Dev Biol. 89:137–162.

Bahmanyar, S., D.D. Kaplan, J.G. Deluca, T.H. Giddings, Jr., E.T. O’Toole, M. Winey, E.D. Salmon, P.J. Casey, W.J. Nelson, and A.I. Barth. 2008. beta-Catenin is a Nek2 substrate involved in centrosome separation. Genes Dev. 22:91–105.

Baskaran, Y., F.P. Tay, E.Y.W. Ng, C.L.F. Swa, S. Wee, J. Gunaratne, and E. Manser. 2021. Proximity proteomics identifies PAK4 as a component of Afadin-Nectin junctions. Nat Commun. 12:5315.

Bauer, D.E., M.C. Canver, and S.H. Orkin. 2015. Generation of genomic deletions in mammalian cell lines via CRISPR/Cas9. JoVE (Journal of Visualized Experiments):e52118.

Bertocchi, C., Y. Wang, A. Ravasio, Y. Hara, Y. Wu, T. Sailov, M.A. Baird, M.W. Davidson, R. Zaidel-Bar, Y. Toyama, B. Ladoux, R.M. Mege, and P. Kanchanawong. 2017. Nanoscale architecture of cadherin-based cell adhesions. Nature cell biology. 19:28–37.

Beutel, O., R. Maraspini, K. Pombo-Garcia, C. Martin-Lemaitre, and A. Honigmann. 2019. Phase Separation of Zonula Occludens Proteins Drives Formation of Tight Junctions. Cell. 179:923–936 e911.

Brasch, J., P.S. Katsamba, O.J. Harrison, G. Ahlsen, R.B. Troyanovsky, I. Indra, A. Kaczynska, B. Kaeser, S. Troyanovsky, B. Honig, and L. Shapiro. 2018. Homophilic and Heterophilic Interactions of Type II Cadherins Identify Specificity Groups Underlying Cell-Adhesive Behavior. Cell Rep. 23:1840–1852.

Bryant, D.M., A. Datta, A.E. Rodriguez-Fraticelli, J. Peranen, F. Martin-Belmonte, and K.E. Mostov. 2010. A molecular network for de novo generation of the apical surface and lumen. Nature cell biology. 12:1035–1045.

Buckley, C.D., J. Tan, K.L. Anderson, D. Hanein, N. Volkmann, W.I. Weis, W.J. Nelson, and A.R. Dunn. 2014. Cell adhesion. The minimal cadherin-catenin complex binds to actin filaments under force. Science (New York, N.Y. 346:1254211.

Buckley, C.E., and D. St Johnston. 2022. Apical-basal polarity and the control of epithelial form and function. Nature reviews. 23:559–577.

Carminati, M., S. Gallini, L. Pirovano, A. Alfieri, S. Bisi, and M. Mapelli. 2016. Concomitant binding of Afadin to LGN and F-actin directs planar spindle orientation. Nat Struct Mol Biol. 23:155–163.

Charras, G., and A.S. Yap. 2018. Tensile forces and mechanotransduction at cell–cell junctions. Current Biology. 28:R445–R457.

Chetty, A.K., J.A. Sexton, B.H. Ha, B.E. Turk, and T.J. Boggon. 2020. Recognition of physiological phosphorylation sites by p21-activated kinase 4. J Struct Biol. 211:107553.

Chiu, C.W., C. Monat, M. Robitaille, M. Lacomme, A.M. Daulat, G. Macleod, H. McNeill, M. Cayouette, and S. Angers. 2016. SAPCD2 controls spindle orientation and asymmetric divisions by negatively regulating the Gαi-LGN-NuMA ternary complex. Developmental cell. 36:50–62.

Choi, H.J., J.C. Gross, S. Pokutta, and W.I. Weis. 2009. Interactions of plakoglobin and beta-catenin with desmosomal cadherins: basis of selective exclusion of alpha- and beta-catenin from desmosomes. J Biol Chem. 284:31776–31788.

Chu, X., X. Chen, Q. Wan, Z. Zheng, and Q. Du. 2016. Nuclear Mitotic Apparatus (NuMA) Interacts with and Regulates Astrin at the Mitotic Spindle. The Journal of biological chemistry. 291:20055–20067.

Clevers, H. 2006. Wnt/β-catenin signaling in development and disease. Cell. 127:469–480.

Culurgioni, S., A. Alfieri, V. Pendolino, F. Laddomada, and M. Mapelli. 2011. Inscuteable and NuMA proteins bind competitively to Leu-Gly-Asn repeat-enriched protein (LGN) during asymmetric cell divisions. Proceedings of the National Academy of Sciences. 108:20998–21003.

di Pietro, F., A. Echard, and X. Morin. 2016. Regulation of mitotic spindle orientation: an integrated view. EMBO reports. 17:1106–1130.

Dixit, R., B. Barnett, J.E. Lazarus, M. Tokito, Y.E. Goldman, and E.L. Holzbaur. 2009. Microtubule plus-end tracking by CLIP-170 requires EB1. Proceedings of the National Academy of Sciences of the United States of America. 106:492–497.

Donà, F., S. Eli, and M. Mapelli. 2022. Insights into mechanisms of oriented division from studies in 3D cellular models. Frontiers in Cell and Developmental Biology. 10:847801.

Du, Q., and I.G. Macara. 2004. Mammalian Pins is a conformational switch that links NuMA to heterotrimeric G proteins. Cell. 119:503–516.

Durgan, J., N. Kaji, D. Jin, and A. Hall. 2011. Par6B and atypical PKC regulate mitotic spindle orientation during epithelial morphogenesis. The Journal of biological chemistry. 286:12461–12474.

Eli, S., G. Rauso, P. Ghezzi, J.L.A. Szczerkowski, M. Bruzzi, F. Rizzelli, F. Iommazzo, A. Loffreda, F. Castagna, F. Dona, C. Gaddoni, A. Dondi, M. Marenda, S. Rodighiero, P. Tournier, Z. Lavagnino, D. Parazzoli, N.C. Gauthier, S. Tamburri, D. Pasini, S.J. Habib, and M. Mapelli. 2025. Localized Wnt-signaling promotes asymmetric NuMA-dependent oriented divisions and unequal apportioning of mitochondria. Nat Commun. 16:10690.

Fankhaenel, M., F.S.G. Hashemi, L. Mourao, E. Lucas, M.M. Hosawi, P. Skipp, X. Morin, C.L. Scheele, and S. Elias. 2023. Annexin A1 is a polarity cue that directs mitotic spindle orientation during mammalian epithelial morphogenesis. Nat Commun. 14:151.

Fujiwara, Y., N. Goda, T. Tamashiro, H. Narita, K. Satomura, T. Tenno, A. Nakagawa, M. Oda, M. Suzuki, T. Sakisaka, Y. Takai, and H. Hiroaki. 2015. Crystal structure of afadin PDZ domain-nectin-3 complex shows the structural plasticity of the ligand-binding site. Protein Sci. 24:376–385.

Fumoto, K., M. Kadono, N. Izumi, and A. Kikuchi. 2009. Axin localizes to the centrosome and is involved in microtubule nucleation. EMBO Rep. 10:606–613.

Gloerich, M., J.M. Bianchini, K.A. Siemers, D.J. Cohen, and W.J. Nelson. 2017. Cell division orientation is coupled to cell-cell adhesion by the E-cadherin/LGN complex. Nat Commun. 8:13996.

Golub, O., B. Wee, R.A. Newman, N.M. Paterson, and K.E. Prehoda. 2017. Activation of Discs large by aPKC aligns the mitotic spindle to the polarity axis during asymmetric cell division. Elife. 6.

Gong, R., M.J. Reynolds, X. Sun, and G.M. Alushin. 2025. Afadin mediates cadherin-catenin complex clustering on F-actin linked to cooperative binding and filament curvature. Sci Adv. 11:eadu0989.

Hadjihannas, M.V., M. Bruckner, and J. Behrens. 2010. Conductin/axin2 and Wnt signalling regulates centrosome cohesion. EMBO Rep. 11:317–324.

Hao, Y., Q. Du, X. Chen, Z. Zheng, J.L. Balsbaugh, S. Maitra, J. Shabanowitz, D.F. Hunt, and I.G. Macara. 2010. Par3 controls epithelial spindle orientation by aPKC-mediated phosphorylation of apical Pins. Curr Biol. 20:1809–1818.

Hart, K.C., J. Tan, K.A. Siemers, J.Y. Sim, B.L. Pruitt, W.J. Nelson, and M. Gloerich. 2017. E-cadherin and LGN align epithelial cell divisions with tissue tension independently of cell shape. Proceedings of the National Academy of Sciences. 114:E5845–E5853.

He, Y.T., L.L. Yang, S.M. Luo, W. Shen, S. Yin, and Q.Y. Sun. 2019. PAK4 Regulates Actin and Microtubule Dynamics during Meiotic Maturation in Mouse Oocyte. Int J Biol Sci. 15:2408–2418.

Huber, A.H., and W.I. Weis. 2001. The structure of the beta-catenin/E-cadherin complex and the molecular basis of diverse ligand recognition by beta-catenin. Cell. 105:391–402.

Hueschen, C.L., S.J. Kenny, K. Xu, and S. Dumont. 2017. NuMA recruits dynein activity to microtubule minus-ends at mitosis. Elife. 6.

Humphries, J.D., A. Byron, M.D. Bass, S.E. Craig, J.W. Pinney, D. Knight, and M.J. Humphries. 2009. Proteomic analysis of integrin-associated complexes identifies RCC2 as a dual regulator of Rac1 and Arf6. Science signaling. 2:ra51.

Ikeda, W., H. Nakanishi, J. Miyoshi, K. Mandai, H. Ishizaki, M. Tanaka, A. Togawa, K. Takahashi, H. Nishioka, H. Yoshida, A. Mizoguchi, S. Nishikawa, and Y. Takai. 1999. Afadin: A key molecule essential for structural organization of cell-cell junctions of polarized epithelia during embryogenesis. The Journal of cell biology. 146:1117–1132.

Itoh, M., A. Nagafuchi, S. Moroi, and S. Tsukita. 1997. Involvement of ZO-1 in cadherin-based cell adhesion through its direct binding to alpha catenin and actin filaments. The Journal of cell biology. 138:181–192.

Kaplan, D.D., T.E. Meigs, P. Kelly, and P.J. Casey. 2004. Identification of a role for β-catenin in the establishment of a bipolar mitotic spindle. Journal of biological chemistry. 279:10829–10832.

Kaushik, R., F. Yu, W. Chia, X. Yang, and S. Bahri. 2003. Subcellular localization of LGN during mitosis: evidence for its cortical localization in mitotic cell culture systems and its requirement for normal cell cycle progression. Mol Biol Cell. 14:3144–3155.

Kiyomitsu, T. 2019. The cortical force-generating machinery: how cortical spindle-pulling forces are generated. Curr Opin Cell Biol. 60:1–8.

Kiyomitsu, T., and S. Boerner. 2021. The nuclear mitotic apparatus (NuMA) protein: A key player for nuclear formation, spindle assembly, and spindle positioning. Frontiers in cell and developmental biology. 9:653801.

Kletter, T., S. Reusch, T. Cavazza, N. Dempewolf, C. Tischer, and S. Reber. 2022. Volumetric morphometry reveals spindle width as the best predictor of mammalian spindle scaling. The Journal of cell biology. 221.

Koelman, E.M.R., A. Yeste-Vazquez, and T.N. Grossmann. 2022. Targeting the interaction of beta-catenin and TCF/LEF transcription factors to inhibit oncogenic Wnt signaling. Bioorg Med Chem. 70:116920.

Kotak, S., C. Busso, and P. Gonczy. 2012. Cortical dynein is critical for proper spindle positioning in human cells. The Journal of cell biology. 199:97–110.

Kuno, S., R. Nakamura, T. Otani, and H. Togashi. 2025. Multivalent afadin interaction promotes IDR-mediated condensate formation and junctional separation of epithelial cells. Cell Rep. 44.

Laan, L., N. Pavin, J. Husson, G. Romet-Lemonne, M. van Duijn, M.P. Lopez, R.D. Vale, F. Julicher, S.L. Reck-Peterson, and M. Dogterom. 2012. Cortical dynein controls microtubule dynamics to generate pulling forces that position microtubule asters. Cell. 148:502–514.

Le, S., M. Yu, and J. Yan. 2019. Phosphorylation Reduces the Mechanical Stability of the alpha-Catenin/ beta-Catenin Complex. Angewandte Chemie (International ed. 58:18663–18669.

Lechler, T., and M. Mapelli. 2021. Spindle positioning and its impact on vertebrate tissue architecture and cell fate. Nature Reviews Molecular Cell Biology. 22:691–708.

Li, Y., Y. Shao, Y. Tong, T. Shen, J. Zhang, Y. Li, H. Gu, and F. Li. 2012. Nucleo-cytoplasmic shuttling of PAK4 modulates beta-catenin intracellular translocation and signaling. Biochim Biophys Acta. 1823:465–475.

Lough, K.J., K.M. Byrd, C.P. Descovich, D.C. Spitzer, A.J. Bergman, G.M. Beaudoin, 3rd, L.F. Reichardt, and S.E. Williams. 2019. Telophase correction refines division orientation in stratified epithelia. Elife. 8.

Lujan, P., T. Rubio, G. Varsano, and M. Kohn. 2017. Keep it on the edge: The post-mitotic midbody as a polarity signal unit. Commun Integr Biol. 10:e1338990.

Martin-Belmonte, F., A. Gassama, A. Datta, W. Yu, U. Rescher, V. Gerke, and K. Mostov. 2007. PTEN-mediated apical segregation of phosphoinositides controls epithelial morphogenesis through Cdc42. Cell. 128:383–397.

Martin-Belmonte, F., W. Yu, A.E. Rodriguez-Fraticelli, A.J. Ewald, Z. Werb, M.A. Alonso, and K. Mostov. 2008. Cell-polarity dynamics controls the mechanism of lumen formation in epithelial morphogenesis. Curr Biol. 18:507–513.

Mirdita, M., K. Schutze, Y. Moriwaki, L. Heo, S. Ovchinnikov, and M. Steinegger. 2022. ColabFold: making protein folding accessible to all. Nat Methods. 19:679–682.

Monster, J.L., C. Manzato, J.A. Van Der Beek, W.-J. Pannekoek, J.A. Hummelink, M.A. Hadders, C. De Heus, J. Klumperman, J. Schuijers, and M. Gloerich. 2025. β-catenin condensation facilitates clustering of the cadherin/catenin complex and formation of nascent cell-cell junctions. Nat Commun.

Mora-Bermudez, F., F. Matsuzaki, and W.B. Huttner. 2014. Specific polar subpopulations of astral microtubules control spindle orientation and symmetric neural stem cell division. Elife. 3.

Morales-Camilo, N., J. Liu, M.J. Ramírez, P. Canales-Salgado, J.J. Alegría, X. Liu, H.T. Ong, N.P. Barrera, A. Fierro, and Y. Toyama. 2024. Alternative molecular mechanisms for force transmission at adherens junctions via β-catenin-vinculin interaction. Nat Commun. 15:5608.

Murray, B.W., C. Guo, J. Piraino, J.K. Westwick, C. Zhang, J. Lamerdin, E. Dagostino, D. Knighton, C.M. Loi, M. Zager, E. Kraynov, I. Popoff, J.G. Christensen, R. Martinez, S.E. Kephart, J. Marakovits, S. Karlicek, S. Bergqvist, and T. Smeal. 2010. Small-molecule p21-activated kinase inhibitor PF-3758309 is a potent inhibitor of oncogenic signaling and tumor growth. Proceedings of the National Academy of Sciences of the United States of America. 107:9446–9451.

Nanes, B.A., C. Chiasson-MacKenzie, A.M. Lowery, N. Ishiyama, V. Faundez, M. Ikura, P.A. Vincent, and A.P. Kowalczyk. 2012. p120-catenin binding masks an endocytic signal conserved in classical cadherins. The Journal of cell biology. 199:365–380.

Nusse, R., and H. Clevers. 2017. Wnt/beta-Catenin Signaling, Disease, and Emerging Therapeutic Modalities. Cell. 169:985–999.

Ooshio, T., R. Kobayashi, W. Ikeda, M. Miyata, Y. Fukumoto, N. Matsuzawa, H. Ogita, and Y. Takai. 2010. Involvement of the interaction of afadin with ZO-1 in the formation of tight junctions in Madin-Darby canine kidney cells. The Journal of biological chemistry. 285:5003–5012.

Otani, T., T.P. Nguyen, S. Tokuda, K. Sugihara, T. Sugawara, K. Furuse, T. Miura, K. Ebnet, and M. Furuse. 2019. Claudins and JAM-A coordinately regulate tight junction formation and epithelial polarity. Journal of Cell Biology. 218:3372–3396.

Ouyang, M., J.Y. Yu, Y. Chen, L. Deng, and C.L. Guo. 2021. Cell-extracellular matrix interactions in the fluidic phase direct the topology and polarity of self-organized epithelial structures. Cell Proliferation. 54:e13014.

Ozawa, M., H. Baribault, and R. Kemler. 1989. The cytoplasmic domain of the cell adhesion molecule uvomorulin associates with three independent proteins structurally related in different species. Embo J. 8:1711–1717.

Pannekoek, W.J., J. de Rooij, and M. Gloerich. 2019. Force transduction by cadherin adhesions in morphogenesis. F1000Res. 8.

Pirovano, L., S. Culurgioni, M. Carminati, A. Alfieri, S. Monzani, V. Cecatiello, C. Gaddoni, F. Rizzelli, J. Foadi, and S. Pasqualato. 2019. Hexameric NuMA: LGN structures promote multivalent interactions required for planar epithelial divisions. Nat Commun. 10:2208.

Pokutta, S., and W.I. Weis. 2000. Structure of the dimerization and beta-catenin-binding region of alpha-catenin. Mol Cell. 5:533–543.

Rajasekaran, A.K., M. Hojo, T. Huima, and E. Rodriguez-Boulan. 1996. Catenins and zonula occludens-1 form a complex during early stages in the assembly of tight junctions. The Journal of cell biology. 132:451–463.

Rizzelli, F., M.G. Malabarba, S. Sigismund, and M. Mapelli. 2020. The crosstalk between microtubules, actin and membranes shapes cell division. Open Biol. 10:190314.

Saadaoui, M., M. Machicoane, F. Di Pietro, F. Etoc, A. Echard, and X. Morin. 2014. Dlg1 controls planar spindle orientation in the neuroepithelium through direct interaction with LGN. Journal of Cell Biology. 206:707–717.

Saw, T.B., X. Gao, M. Li, J. He, A.P. Le, S. Marsh, K.-h. Lin, A. Ludwig, J. Prost, and C.T.J.N.P. Lim. 2022. Transepithelial potential difference governs epithelial homeostasis by electromechanics.1–7.

Selamat, W., P.L. Tay, Y. Baskaran, and E. Manser. 2015. The Cdc42 Effector Kinase PAK4 Localizes to Cell-Cell Junctions and Contributes to Establishing Cell Polarity. PLoS One. 10:e0129634.

Seldin, L., N.D. Poulson, H.P. Foote, and T. Lechler. 2013. NuMA localization, stability, and function in spindle orientation involve 4.1 and Cdk1 interactions. Mol Biol Cell. 24:3651–3662.

Solanas, G., S. Miravet, D. Casagolda, J. Castano, I. Raurell, A. Corrionero, A. Garcia de Herreros, and M. Dunach. 2016. beta-Catenin and plakoglobin N- and C-tails determine ligand specificity. The Journal of biological chemistry. 291:23925–23927.

Speicher, S., A. Fischer, J. Knoblich, and A. Carmena. 2008. The PDZ protein Canoe regulates the asymmetric division of Drosophila neuroblasts and muscle progenitors. Curr Biol. 18:831–837.

Sun, M., M. Jia, H. Ren, B. Yang, W. Chi, G. Xin, Q. Jiang, and C. Zhang. 2021. NuMA regulates mitotic spindle assembly, structural dynamics and function via phase separation. Nat Commun. 12:7157.

Takayanagi, H., J. Hayase, S. Kamakura, K. Miyano, K. Chishiki, S. Yuzawa, and H. Sumimoto. 2019. Intramolecular interaction in LGN, an adaptor protein that regulates mitotic spindle orientation. The Journal of biological chemistry. 294:19655–19666.

Takeichi, M. 2014. Dynamic contacts: rearranging adherens junctions to drive epithelial remodelling. Nature Reviews Molecular Cell Biology. 15:397–410.

Tame, M.A., J.A. Raaijmakers, P. Afanasyev, and R.H. Medema. 2016. Chromosome misalignments induce spindle-positioning defects. EMBO Rep. 17:317–325.

Tang, Z., Y. Hu, Z. Wang, K. Jiang, C. Zhan, W.F. Marshall, and N. Tang. 2018. Mechanical Forces Program the Orientation of Cell Division during Airway Tube Morphogenesis. Dev Cell. 44:313–325 e315.

Tuncay, H., B.F. Brinkmann, T. Steinbacher, A. Schurmann, V. Gerke, S. Iden, and K. Ebnet. 2015. JAM-A regulates cortical dynein localization through Cdc42 to control planar spindle orientation during mitosis. Nat Commun. 6:8128.

Valenta, T., G. Hausmann, and K. Basler. 2012. The many faces and functions of beta-catenin. Embo J. 31:2714–2736.

Vasioukhin, V., C. Bauer, L. Degenstein, B. Wise, and E. Fuchs. 2001. Hyperproliferation and defects in epithelial polarity upon conditional ablation of alpha-catenin in skin. Cell. 104:605–617.

Wang, X., B. Dong, K. Zhang, Z. Ji, C. Cheng, H. Zhao, Y. Sheng, X. Li, L. Fan, and W. Xue. 2018. E-cadherin bridges cell polarity and spindle orientation to ensure prostate epithelial integrity and prevent carcinogenesis in vivo. PLoS Genet. 14:e1007609.

Wee, B., C.A. Johnston, K.E. Prehoda, and C.Q. Doe. 2011. Canoe binds RanGTP to promote Pins(TPR)/Mud-mediated spindle orientation. The Journal of cell biology. 195:369–376.

Wu, J., P. Rowart, F. Jouret, B.M. Gassaway, V. Rajendran, J. Rinehart, and M.J. Caplan. 2020. Mechanisms involved in AMPK-mediated deposition of tight junction components to the plasma membrane. Am J Physiol Cell Physiol. 318:C486–C501.

Yonemura, S., Y. Wada, T. Watanabe, A. Nagafuchi, and M. Shibata. 2010. alpha-Catenin as a tension transducer that induces adherens junction development. Nature cell biology. 12:533–542.

Yu, H.H., M.R. Dohn, N.O. Markham, R.J. Coffey, and A.B. Reynolds. 2016. p120-catenin controls contractility along the vertical axis of epithelial lateral membranes. Journal of cell science. 129:80–94.

Yuzawa, S., S. Kamakura, Y. Iwakiri, J. Hayase, and H. Sumimoto. 2011. Structural basis for interaction between the conserved cell polarity proteins Inscuteable and Leu-Gly-Asn repeat-enriched protein (LGN). Proceedings of the National Academy of Sciences of the United States of America. 108:19210–19215.

Zhan, L., A. Rosenberg, K.C. Bergami, M. Yu, Z. Xuan, A.B. Jaffe, C. Allred, and S.K. Muthuswamy. 2008. Deregulation of scribble promotes mammary tumorigenesis and reveals a role for cell polarity in carcinoma. Cell. 135:865–878.

Zheng, Z., Q. Wan, G. Meixiong, and Q. Du. 2014. Cell cycle-regulated membrane binding of NuMA contributes to efficient anaphase chromosome separation. Mol Biol Cell. 25:606–619.

Zheng, Z., H. Zhu, Q. Wan, J. Liu, Z. Xiao, D.P. Siderovski, and Q. Du. 2010. LGN regulates mitotic spindle orientation during epithelial morphogenesis. The Journal of cell biology. 189:275–288.

Zhu, J., W. Wen, Z. Zheng, Y. Shang, Z. Wei, Z. Xiao, Z. Pan, Q. Du, W. Wang, and M. Zhang. 2011. LGN/mInsc and LGN/NuMA complex structures suggest distinct functions in asymmetric cell division for the Par3/mInsc/LGN and Gαi/LGN/NuMA pathways. Mol Cell. 43:418–431.

